# The endolysosomal phospholipid bis(monoacylglycero)phosphate is synthesized via intra- and extracellular pathways

**DOI:** 10.1101/2024.05.28.596187

**Authors:** Dominik Bulfon, Johannes Breithofer, Gernot F. Grabner, Nermeen Fawzy, Anita Pirchheim, Heimo Wolinski, Dagmar Kolb, Lennart Hartig, Martin Tischitz, Clara Zitta, Greta Bramerdorfer, Achim Lass, Ulrike Taschler, Dagmar Kratky, Peter Greimel, Robert Zimmermann

## Abstract

Bis(monoacylglycero)phosphate (BMP) is a major phospholipid constituent of intralumenal membranes in late endosomes/lysosomes, where it regulates the degradation and sorting of lipid cargo. Recent observations suggest that the Batten disease - associated protein CLN5 functions as lysosomal BMP synthase. Here, we show that transacylation reactions catalyzed by cytosolic and secreted enzymes enhance BMP synthesis independently of CLN5. The transacylases identified in this study are capable of acylating the precursor lipid phosphatidylglycerol (PG), generating acyl-PG, which is subsequently hydrolyzed to BMP. Extracellularly, acyl-PG and BMP are generated by endothelial lipase in cooperation with other serum enzymes of the pancreatic lipase family. The intracellular acylation of PG is catalyzed by several members of the cytosolic phospholipase A2 group IV (PLA2G4) family. Overexpression of secreted or cytosolic transacylases was sufficient to correct BMP deficiency in HEK293 cells lacking *CLN5*. Collectively, our observations suggest that functionally overlapping pathways promote BMP synthesis in mammalian cells.

## Introduction

Cells take up nutrients, ligands, and components of the plasma membrane via the endocytic pathway. This pathway is essential for maintaining metabolic homeostasis and composed of organelles that undergo dynamic transformation, comprising early endosomes, multivesicular bodies, late endosomes (LE), and lysosomes. During maturation, the inward budding of the limiting endosomal membrane produces intralumenal vesicles (ILVs), which results in the formation of multivesicular organelles. This process is essential for organelle maturation and proper cargo sorting^1^. ILVs undergo remodeling during organelle maturation leading to substantial changes in lipid composition^2^. Bis(monoacylglycero)phosphate (BMP), also known as lysobisphosphatidic acid (LBPA), is a major lipid constituent of internal membranes of LE and lysosomes^2^. This phospholipid interacts with many lumenal proteins and thereby promotes the degradation and export of lipid cargo^3,4^.

The availability of lumenal membranes in lysosomes is particularly important for lipid hydrolases, since these enzymes require a water-lipid interphase for activation^5^. BMP is negatively charged and highly resistant to degradation in the acidic lysosomal compartments. Based on these features, it can form a docking station for lumenal hydrolases that are positively charged at acidic pH^5^. Accordingly, it has been shown that BMP stimulates the activity of lysosomal lipid hydrolases catalyzing the degradation of glycerophospholipids, sphingolipids, and neutral lipids^5,6^.

BMP also interacts with lipid-binding and -transport proteins. It supports the presentation of glycolipids to catabolic enzymes by sphingolipid activator proteins^7,8^. Furthermore, BMP facilitates cholesterol export from lysosomes^9,10^, which depends on Niemann-Pick type C1 and C2 proteins (NPC1 and NPC2). NPC2 directly interacts with BMP and this interaction strongly promotes cholesterol exchange between membranes^11^. Recent reports demonstrated that BMP supplementation and drug-induced stimulation of BMP synthesis can reduce cholesterol overload in NPC1-deficient cells and mice ^12,13^.

Under pathological conditions, BMP accumulates in numerous lysosomal storage disorders caused by mutations in genes encoding lysosomal proteins^14^ or as a side effect of cationic amphiphilic drug induced lipidosis^15^. Furthermore, altered BMP levels are lipidomic signatures of common neurodegenerative diseases such as Parkinson’s ^16^ and Alzheimer’s disease^17,18^. BMP accumulation occurs in response to pathological lipid accumulation and facilitates cargo degradation and sorting^19^. Conversely, a reduced cellular BMP content may compromise lysosomal lipid degradation and sorting pathways. Recent observations suggest that BMP deficiency precedes and may underlie the development of lipidosis in cells lacking progranulin (PGRN)^20,21^ and ceroid lipfuscinosis neuronal protein 5 (CLN5)^22^. Loss-of-function mutations in these genes are causal for two of the thirteen forms of Batten disease (CLN11 and CLN5), a rare neurodegenerative disease characterized by the rapid deterioration of cognition and movement^23^. PGRN and CLN5 affect BMP synthesis by different mechanisms. CLN5 functions as a lysosomal transacylase that generates BMP by acylating lyso-phosphatidylglycerol^22^. PGRN interacts directly with BMP and mice lacking PGRN show global BMP deficiency. It is currently unclear whether the BMP/PGRN interaction modulates BMP stability or the activity of BMP-metabolizing enzymes^20,21^.

BMP has also been implicated in the pathogenesis of antiphospholipid syndrome (APS), an autoimmune disease associated with arterial and venous thrombosis and pregnancy-related complications^24^. The hypercoagulable state is caused by high levels of antiphospholipid antibodies, which activate coagulation pathways and induce inflammation. These antibodies can directly bind anionic phospholipids such as cardiolipin and BMP, or target proteins that bind anionic phospholipids^25^. A recent study identified BMP in complex with endothelial protein C receptor as a pathogenic cell surface antigen that interacts with antiphospholipid antibodies. The authors proposed that this interaction represents a central mechanism for the development and progression of autoimmune diseases^26^.

ILVs and BMP can influence viral infection, as viruses hijack the endosomal machinery for cell entry. Viral entry involves clathrin-mediated endocytosis and fusion of viral and endosomal membranes. The viral envelope can fuse directly with the limiting membrane of endocytic vesicles, leading to the release of the viral capsid into the cytosol. Alternatively, the virus undergoes fusion with ILVs, resulting in the release of the viral capsid into the protective environment of the ILV lumen^27^. The capsid is then exported via backfusion of the ILVs with the limiting membrane. In some cases, the release of the capsid into the cytosol depends on BMP and the BMP-binding protein ALG-2-interacting protein X (ALIX)^27^. BMP was identified as co-factor for several viruses^3^ including influenza virus^28^, and was also proposed to affect infections with the novel coronavirus SARS-CoV-2^29^.

Although numerous studies suggested that BMP is highly relevant for metabolic homeostasis and human disease, fundamental aspects of BMP metabolism are incompletely understood, including the molecular mechanisms mediating its biosynthesis and degradation. Currently, the prevailing view is that BMP is synthesized in acidic organelles from the precursor lipid phosphatidylglycerol (PG), as extensively reviewed by Hullin-Matsuda *et al.*^4^, and CLN5 was identified as lysosomal BMP synthase^22^. However, several *in vitro* studies indicate that other mechanisms contribute to this process, such as PG-independent BMP formation^30,31^ or BMP formation in non-acidic compartments^32^. Here, we show that secreted and cytosolic transacylases of the pancreatic lipase (PNLIP) and the phospholipase A2 group IV (PLA2G4) enzyme family promote BMP synthesis in mammalian cells.

## Methods

### Materials

Lipids and internal standards were purchased from Avanti Polar Lipids. For storage, the lipid stock solutions were dissolved in CHCl_3_, covered with a layer of N_2_, and kept at −20°C. All other materials and consumables were purchased form Sigma Aldrich, unless indicated otherwise.

### Cell culture

HEK293 cells, COS-7 cells, and HepG2 cells were grown in DMEM (4.5 g/l glucose; Gibco, Cat#11965084) containing 10% fetal bovine serum (FBS), 100 U/mL penicillin, and 100 µg/mL streptomycin (P/S) under standard conditions (95% humidified atmosphere, 37°C, 7% CO_2_). All cell culture experiments were conducted using cells below a passage number of 30. Expi293F cells (derived from HEK293 cells) were grown in suspension and shaken at 125 rpm using serum-free Expi293^TM^ expression medium (Gibco, Cat#A1435101) according to the manufacturer’s instructions.

### Knockout cell lines

*PLA2G4B*, *PLA2G4C*, and *CLN5* were deleted in HEK293 cells by CRISPR/Cas9 using the pSpCas9(BB)-2A-Puro (PX459) V2.0 vector (Addgene, plasmid #62988) and two to three sgRNAs per gene (Table S1), designed with [chopchop](https://chopchop.cbu.uib.no/)^33^. sgRNA were cloned into the *BbsI* site of pSPCas9(BB)-2A-Puro V2.0. Correct insertion was verified by Sanger sequencing. The resulting constructs were transfected into HEK293 cells using the Lipofectamine 3000 Reagent (Invitrogen, Cat#L3000001) according to the manufacturer’s instructions. Cells transfected with empty pSPCas9(BB)-2A-Puro V2.0 served as control. Twenty-four hours after transfection, 2 µg/ml puromycin were added for selection of successfully transfected cells. After clonal expansion for three weeks, the cell lines were screened for PLA2G4B, PLA2G4C, and CLN5 deletion by Sanger sequencing using primers (Table S1) that bind the region with the expected frameshift. Control cells transfected with an empty vector were subjected to the same procedure, including passaging and validation of the genomic sequence by Sanger sequencing.

### Animals

Mice were maintained on a regular light-dark cycle (14 h light, 10 h dark) at a room temperature (RT) of 23 ± 1°C and kept *ad libitum* on a standard laboratory chow diet (4.5% w/w fat, Ssniff Spezialdiaeten, R/M-H Extrudate, V1126-027). Male mice aged 15–21 weeks were used for the experiments. B6.129S-Lipg^tm1Tq/J^ (*Lipg*-ko) mice were obtained from the Jackson Laboratory (Bar Harbor, ME). Animals were anesthetized with IsoFlo/Isoflurane (Abbott, Animal Health, Queenborough, Kent, UK) and sacrificed by cervical dislocation. All studies involving animals are reported in accordance with the ARRIVE guidelines for reporting experiments involving animals^34^. Experimental procedures were approved by the Ethics committee of the University of Graz, and the Austrian Federal Ministry of Education, Science and Research (protocol number 2022-0.466.992) and were conducted in accordance with the council of Europe Convention (ETS 123). All animal procedures were performed as humanely as possible to minimize suffering.

### Synthesis of stably labeled phosphatidylglycerol

Phosphatidylglycerol (PG) containing uniformly labelled (UL)-^13^C-glycerol as head group (^13^C-PG) was synthesized by the conversion of di-oleoyl phosphatidylcholine (PC) into di-oleoyl PG by cabbage phospholipase D (PLD) as described previously^35^. Briefly, PLD (Sigma Aldrich, Cat#P8398) was dissolved in 1 ml 0.2 M acetate buffer (pH 5.6) containing 40 mM CaCI_2_ and 30% ^13^C-glycerol (w/w) (Sigma-Aldrich, Cat#489476). Forty micrograms of 1,2-dioleoyl-PC were dissolved in 1.7 ml diethyl ether. One ml of the aqueous enzyme solution was incubated with 1.7 ml PC-ether solution at 30°C under continuous stirring (650 rpm) for 4h. Subsequently, lipids were extracted by washing the aqueous phase three times with 1.7 ml diethyl ether. The lipid extract was dried under a stream of N_2_ and then dissolved in 0.5 ml CHCl_3_. PG was isolated by solid phase extraction using a Strata^®^ NH_2_ column (Phenomenex, Cat#8B-S009-JCL). For this purpose, the NH_2_ column was equilibrated with 4 ml hexane and lipids were applied. PC was eluted with 4 ml CHCl_3_/MeOH/H_2_O (3:7:1 v/v), followed by a wash with 4 ml diethyl ether, 1% acetic acid. PG was eluted with 6 ml CHCl_3_ /MeOH/0.8 M ammonium acetate (3:7:1, v/v/v). Phase separation was induced by the addition of 6 ml CHCl_3_ and 2 ml ddH_2_O. The organic phase was collected and dried under a stream of N_2_. ^13^C-PG was analyzed and quantitated using LC-MS.

### Lipid Extraction

Lipid samples from *in vitro* and *in vivo* experiments were analyzed by LC-MS. For cell-free assays, lipids were extracted according to Matyash et al.^36^ with modifications, 1 ml methyl-tert-butylether (MTBE)/MeOH (3:1; v/v) containing 500 pmol butylated hydroxytoluene (BHT), 0.01% acetic acid, and 150 pmol internal standard (14:0-14:0 BMP, 14:0-14:0-14:0 hemi-BMP, 17:0-17:0 PG) per sample. Samples were extracted under constant shaking on a thermomixer at 1,200 rpm for 30 min at RT. After the addition of 200 µl ddH_2_O (100 µl for *in vitro* assays) and further incubation for 10 min at RT, samples were centrifuged for phase separation at 20,000 x *g* for 5 min at RT. The upper organic phase was collected, dried under a stream of N_2,_ and reconstituted in MeOH/2-propanol/ddH_2_O (6:3:1; v/v/v) for UPLC/MS analysis.

Samples of cell pellets, medium (70µl), or tissue explants were transferred to 2-ml safe-lock tubes containing two 5-mm steel beads. Lipids were extracted using 700 μl MTBE/MeOH (3:1; v/v) containing 500 pmol BHT, 0.01% acetic acid, and 150 pmol internal standards (14:0-14:0 BMP, 14:0-14:0-14:0 hemi-BMP, 17:0-17:0 PG, 17:1 LPG) per sample. Samples were homogenized by shaking on a Retsch Tissue-Lyser (Qiagen) for 2×10 sec (30 Hz). Lipids were extracted under constant shaking on a thermomixer at 1,200 rpm for 30 min at RT. After the addition of 140 µl (70µl for medium) ddH_2_O and further incubation for 10 min at RT, samples were centrifuged for phase separation at 20,000 x *g* for 5 min at RT. The upper organic phase was collected, dried under a stream of N_2,_ and reconstituted in MeOH/2-propanol/ddH_2_O (6:3:1; v/v/v) for UPLC/MS analysis. Proteins in the lower aqueous phase were dried in the oven and dissolved in 200 µl NaOH/SDS (0.3M/0.1%). The protein concentration was determined using the Pierce BCA Protein Assay Kit (Thermo Fisher Scientific, Cat#23225).

For TLC analysis, lipids (50µl) were extracted according to Folch^37^ using 1 ml CHCl_3_/MeOH (2:1; v/v) for 20 min on an overtop shaker at RT. Phase separation was achieved by the addition of 200 µl ddH_2_O. Samples were centrifuged at 2,600 x *g* for 5 min, and 800 µl of the lower organic phase were transferred to a 1.5 ml tube and dried under a stream of N_2_.

### Enzyme activity assays

The substrates for transacylase and hydrolase activity assays were prepared by evaporating the phospholipid stock solutions under a constant stream of N_2_, followed by sonication in PBS using a SONOPULS ultrasonic homogenizer (Bandelin) with 15-20% amplitude for 3×10s on ice. All phospholipid substrates were esterified with oleic acid.

#### Transacylase activity assays

Neutral transacylase activity of cell lysates was determined using 20 µl cell lysate (40 µg protein) adjusted to 50 µl with 0.1 M bis-tris propane buffer (pH 7.0). The reaction was started by the addition of 50 µl phospholipid substrate. For determination of the pH optimum of the reaction, the buffer system was replaced by 0.1 M citrate buffer (pH 4.5-5.0), 0.1 M MES buffer (pH 5.5-6), and 0.1 M bis-tris propane (pH 6.5-8). The final reaction mixture contained 0.4 mg/ml cell protein, 0.32 mM phospholipid substrate, 1% FA-free BSA, and the indicated reaction buffer. For partially purified enzymes, the final concentrations were 10 µg protein/ml (1 µg/assay), 1 mM substrate, and 1% FA-free BSA. If indicated, oleoyl-CoA (stock solution 2 mM prepared in 10 mM sodium acetate buffer pH 6.0) and CaCl_2_ were added to a final concentration of 100 µM and 2 mM, respectively. Samples were incubated in 2 ml safe-lock Eppendorf tubes in a thermoshaker for 1h at 37°C and 450 rpm. The reaction was stopped by the addition of 1 ml MBTE extraction mix (see lipid extraction) and reaction products were analyzed by LC-MS or TLC.

To monitor transacylase activity of cell supernatants, we used 20 to 70 µl of conditioned, serum-free medium/assay in a final volume of 100 µl. The final reaction concentrations were 0.125 mM substrate, 1% FA-free BSA and 20 mM HEPES (pH 7.4).

To detect transacylase activity in heparin-plasma, we used 30 µl plasma in a final volume of 50 µl. The final concentrations were 60% plasma, 0.5 mM substrate, and 1% FA-free BSA. To collect heparin-plasma, mice were injected with 1 U / g body weight heparin (Heparin Gilvasan 5000 I.E./ml). Blood was collected 15 min after injection. Blood samples were centrifuged for 10 min at 1000 x g and 4°C. The supernatant (heparin-plasma) was collected and stored at −80°C.

Hemi-BMP synthase activity of FBS was also determined under conditions mimicking cell culture experiments. DMEM containing 10% FBS was adjusted to 50 µM PG (final concentration) and incubated at 37°C for 4h. As controls, we used DMEM without FBS, DMEM containing heat-inactivated FBS, and DMEM containing 40 µM orlistat. For heat-inactivation, 10 ml FBS was pre-warmed to 37°C before being placed in a water bath at 56°C for 30min.

#### Hydrolase activity assays

To determine the hydrolytic activities of PLA2G4 enzymes, 20 µl of cell lysates (2 mg/ml) were incubated with 20 µl substrate (BMP or hemi-BMP). The substrate was prepared in PBS and the final reaction mixture contained 1 mg/ml cell protein, 0.5 mM substrate, and 1% FA-BSA. Assays were performed in the presence and absence of 2 mM CaCl_2_ for 30 min at 37°C. The release of non-esterified FAs was measured using the NEFA-HR (2) enzymatic test (FUJIFILM WAKO Chemicals, Cat#434-91795, Cat#436-91995) following the manufacturer’s instructions.

The hydrolysis of hemi-BMP by partially purified PLA2G4 enzymes was determined by incubating the semi-purified enzymes (1µg protein/assay) with the substrate for 30 min at 37°C in a final volume of 40 µl. The substrate was prepared in PBS and final reaction conditions were 50 µg protein/ml, 0.5 mM hemi-BMP, and 2% FA-free BSA. If indicated, CaCl_2_ and CHAPS were added to the reaction mixture (final concentration 2mM and 2.5mM, respectively). Lipids were extracted using the Folch method and analyzed by TLC.

The monitor the hydrolysis of hemi-BMP by PNLIP enzymes, 20 µl conditioned serum-free medium was mixed with 20 µl substrate containing hemi-BMP. The final reaction concentrations were 50% conditioned medium, 0.5 mM substrate, and 1% FA-BSA. Lipids were extracted using the Folch method and analyzed by TLC.

### Cell culture experiments

To induce hemi-BMP and BMP formation, PG was added to the culture medium (DMEM, 4.5 g/l glucose, 10% FBS, P/S) at a final concentration of 50 µM. If indicated, FBS was replaced by heat-inactivated FBS (56°C, 30 min). After incubation for the indicated time period, cells were washed three-times with PBS, collected by scraping, and centrifuged for 3 min at 300 x *g* and 4°C. Cell pellets were stored at −20°C until further analysis. When the conditioned medium of a cell culture experiment was analyzed, an aliquot of the medium was taken prior to harvesting. The medium was centrifuged at 1,000 x *g* for 10min at 4°C to remove cell debris, and the supernatant was stored at −20°C.

The same procedure was used to study the incorporation ^13^C-PG and ^13^C-glucose into BMP. For this purpose, cells were washed with PBS and incubated with standard medium containing 50 µM ^13^C-PG for 18h. Subsequently, lipids were extracted using the MTBE method and analyzed by LC-MS. To study ^13^C-glucose incorporation, we used DMEM without glucose (Gibco, Cat#11966025) supplemented with 4.5 g/l ^13^C-glucose (Sigma, Cat#389374) and unlabeled PG (50 µM).

### Transfection and preparation of cell lysates

All plasmids (vectors and protein names) encoding recombinant lipid hydrolase/transacylases are listed in Supplemental Information Table S2.

The enzyme library, comprising cell lysates and conditioned media, was established using Expi293F cells expressing recombinant proteins. Expi293F cells (3*10^6^ cells/ml) were transfected in 50 ml CELLSTAR reactor tubes (Greiner Bio-One GmbH, Cat#A1435101) with ExpiFectamine293 Transfection Kit (Gibco, Cat#A14524) in Opti-Mem reduced serum medium (Gibco, Cat#31985062) according to the manufacturer’s instructions. Two days after transfection, cells and culture medium were harvested by centrifugation (300 x *g*, 3 min). Cells were washed three times with PBS and pellets were stored at −20°C. The culture medium was centrifuged at 10,000 x *g* for 10 min at 4°C to remove cell debris before storage at −20°C. To improve the stability of recombinant proteins in conditioned medium, sucrose (0.25 M) and HEPES (20 mM, pH 7.4) were added after collecting the supernatants.

HEK293 cells were transfected with Metafectene (Biontex, Cat#T020-5.0) according to manufacturer’s instructions. Briefly, 2*10^6^ HEK293 cells were seeded in 10 ml medium in a 100-mm cell culture dish. 27 µl Metafectene were mixed with 300 µl DMEM (without FBS and P/S) in a 1.5-ml Eppendorf tube. In a separate 1.5-ml tube, 6 µg DNA were mixed with 300 µl DMEM (without FBS and P/S). The contents were pooled, mixed and incubated for 20 min at RT. The DNA-Metafectene mixture was added dropwise to the dish. After 4h of incubation, the medium was replaced by fresh cell culture medium. After 24h, cells were either trypsinized and re-plated for further experiments or harvested and frozen for Western blot analysis and cell lysate preparation. Cells transfected with an empty vector were used as a control.

To prepare lysates, cells were disrupted in lysis solution (0.25 M sucrose, 1 mM EDTA, 1 mM dithiothreitol, 20 μg/ml leupeptin, 2 μg/ml antipain, 1 μg/ml pepstatin, pH 7.0) by sonication using a SONOPULS ultrasonic homogenizer (Bandelin). Nuclei and debris were removed by centrifugation at 1,000 *x* g and 4°C for 10 min. Protein concentrations of lysates were determined using Bio-Rad protein assay (Bio-Rad, Cat#5000006) and BSA as standard. Lysates were stored at −20°C.

### Site-directed mutagenesis

The primers used to generate catalytically inactive mutants by replacing the active serine with an alanine for murine LIPG, PLA2G4E, and PLA2G4D are listed in Table S1. Mutagenesis was performed using the Q5 site-directed mutagenesis kit (NEB, Cat#E0552S) according to the manufacturer’s instructions. Annealing temperatures were set according to the NCBI primer tool. Elongation time was set to 3:30 min (∼20-30 sec/kb). Constructs were transformed into chemically competent *E. coli*. The base-exchange was verified by Sanger sequencing.

### Protein purification

For protein purification, 25 ml Expi293F culture (3*10^6^ cells/ml) were transfected with indicated plasmids. After 48h, cells were washed three times with 15 ml PBS and resuspended in 60 mM sodium-phosphate buffer (pH 7.4) containing 0.1% NP-40. Cell lysates were prepared by sonication and subsequent centrifugation at 1,000 x g, for 10 min, and 4°C to remove the cell debris and nuclear fraction. Cell lysates were adjusted to 50 mM sodium-phosphate, 300 mM NaCl and subsequently diluted to 50 ml with binding buffer (50 mM sodium-phosphate buffer containing 300 mM NaCl) in a 50 ml tube. TALON^®^ metal affinity resin (500 μl, Takara Bio, Cat#635501) was equilibrated by resuspending the beads twice in 10 ml of binding buffer. Subsequently, the beads were transferred to the lysates and the suspension was incubated overnight on an overtop shaker at 4°C. The affinity beads were removed from the lysates by centrifugation at 300 x *g* for 3 min and transferred onto a gravity column. The beads were washed four times with 4 ml binding buffer. Subsequently, the protein was eluted with 50 mM sodium-phosphate buffer (pH 7) containing 300 mM NaCl and 250 mM imidazole in 20×100 μl elution steps. Protein elution was monitored by the determination of protein concentrations in the fractions. Fractions containing protein were pooled and desalted using the Econo-Pac® 10-DG Desalting Columns (Bio-Rad, Cat#732-2010). To this end, the column was equilibrated with 20 ml of storage buffer (50 mM Tris-HCl pH 7.4, 20 µM DTT, 250 mM sucrose) prior to loading of the pooled fractions. After the whole sample entered the column, additional storage buffer (void volume of 3 ml – sample volume) was loaded. Purified proteins were eluted in storage buffer with 1.5-fold the volume of the sample. Protein aliquots were snap frozen in liquid N_2_ and stored at −80°C.

### Immunoblotting

For expression analysis of recombinant proteins, 10-30 µg protein of cell lysates were mixed with SDS-sample buffer and heated for 10min at 95°C prior to loading on the SDS-polyacrylamide gel. Proteins were separated by applying a constant current of 25 mA. Subsequently, proteins were transferred onto a polyvinylidene fluoride (PVDF) transfer membrane (Roth) using a blotting chamber filled with CAPS buffer (10 mM CAPS, 10% MeOH (v/v), pH 11) and a constant current of 200 mA for 90 min. The membranes were blocked for 1h with 10 % blotting grade milk powder in TST (50 mM Tris/HCl, pH 7.4, 0.15 M NaCl, 0.1 % Tween-20). Membranes were incubated for 2h (at RT) or 24h (at 4°C) with anti-6X His-tag antibody (Abcam, Cat#ab18184) or anti-CLN5 antibody (Abcam, Cat#ab170899) in 5% milk powder in TST. Secondary antibodies (see KRT) were prepared in 5% milk powder in TST. Antibody binding was visualized using Clarity Western ECL Substrate and a ChemiDoc Touch Imaging System (both Bio-Rad).

### qPCR analysis

For isolation of total RNA, HEK293 cells were lysed with 1 ml TRIzol™ and incubated at RT for 5 min. Phase separation was achieved by addition of 100 μl 1-bromo-3-chloropropane and centrifugation at 12,000 × *g* and 4 °C for 15 min. The aqueous supernatant was transferred to a fresh Eppendorf tube and total RNA was precipitated by addition of 500 μl 2-propanol and centrifugation at 12,000 × *g* and 4 °C for 10 min. RNA was washed twice with 70% ethanol and centrifugation at 7,500 × *g* and 4 °C for 5 min. DNA digestion was performed with DNase I and RNA was reverse-transcribed into cDNA using LunaScript^TM^ RT SuperMix kit (New England Biolabs, Cat#E3010L). qPCR was performed using the StepOnePlus^TM^ Real-Time PCR System with SYBR green as the detection method. Primers used for gene expression analysis are listed in Table S1. The *36B4* gene was used as housekeeping gene and relative gene expression was normalized to PLA2G4A signal. Samples were analyzed in technical triplicates.

### Intraperitoneal administration of PG and uptake studies in mice

Male *Lipg*-ko mice and wild-type littermates received an intraperitoneal injection of 5 mg PG in 100 µl saline. Before (time point 0) and 1, 4, and 24 h after injection, blood samples were collected by orbital venous sinus bleeding, and EDTA plasma was prepared.

To investigate the uptake, conversion, and tissue distribution of ^13^C-PG, male *Lipg*-ko and WT littermates received an intraperitoneal injection of 200 µg ^13^C-PG mixed in 100 µl saline. Four hours after injection, blood samples were collected and animals were sacrificed by cervical dislocation. Tissues were collected, washed in PBS containing 1mM EDTA, snap-frozen, and stored at −80°C.

### Thin layer chromatography

Dried lipid extracts were reconstituted in 40 µl CHCl_3_ and spotted onto a TLC Silica gel 60 20×20 cm aluminum sheet (Merck, Cat#1.05553.0001). Chromatography was performed with CHCl_3_/MeOH/H_2_O (65:35:5, v/v/v). The silica plate was immersed in charring solution (25% EtOH, 8.5% H_3_PO_4_, 5% v/ CuSO_4_, v/v/w), and lipid bands were visualized at 130°C. Images of the TLC plates were captured with the ChemiDoc Touch Imaging System (Bio-Rad). Densitometric analysis was performed in ImageJ using the raw TIFF files. Data is expressed in arbitrary units (AU). For better visualization, the brightness was adjusted after the densitometric analysis.

### MS analysis

MS method for screening assays: chromatographic separation was performed using an Agilent 1290 Infinity II UHPLC equipped with an Agilent Eclipse XDB-C18 (3.0×20 mm, 1.8 µm, Cat#HEWL926975-302), a flow rate of 0.6 ml/min, an injection volume of 5 μl, a column temperature of 50°C, and a 6 min linear gradient with H_2_O and mobile phase A and 2-propanol as mobile phase B. Both solvents contained 10 mM ammonium acetate, 0.1% formic acid, and 8 µM phosphoric acid. Hemi-BMP was detected on an Agilent 6470 triple-quadrupole mass spectrometer with Agilent Jet Stream ESI (Agilent Technologies) in ESI positive mode, controlled by Agilent MassHunter Acquisition software version 10.1. Positive mode: [MNH_4_]^+^ to [DG-H_2_O]^+^ of the respective DG for hemi-BMP. Unit resolution was used in both MS1 and MS2, the collision energy was set to 25 V for hemi-BMP.

MS method for BMP, PG, and hemi-BMP MS analysis: Chromatographic separation was performed using an Agilent 1290 Infinity II UHPLC equipped with an ACQUITY UPLC BEH C18 Column 2.1×150 mm, 1.7 µm (Waters Corporation, Cat#186002353), a flow rate of 0.2 ml/min, an injection volume of 2 μl, and a column temperature of 50°C. Mobile phase A was MeOH/H_2_O (8/2, v/v) and mobile phase B was 2-propanol/MeOH (8/2, v/v). Both solvents contained 10 mM ammonium acetate, 0.1% formic acid, and 8 µM phosphoric acid. The chromatography consisted of a 30min gradient starting with a linear increase of 50% to 60% mobile phase B over the time course of 13min, followed by a 7min linear gradient to 100% mobile phase B, which was held for 5min before returning to 50% mobile phase B for 5min. BMP and PG species were detected on an Agilent 6470 triple-quadrupole mass spectrometer with Agilent Jet Stream ESI (Agilent Technologies) under fast polarity switching with simultaneous analysis in ESI positive and negative modes in a single run, controlled by Agilent MassHunter Acquisition software version 10.1. Lipid species were analyzed in dynamic multiple reaction monitoring mode using [M-H]^-^ to FA anion transitions in the negative mode. Positive mode: [MNH_4_]^+^ to [MG-H_2_O]^+^ of the respective MG for BMP and [MNH_4_]^+^ to [DG-H_2_O]^+^ of the respective DG for PG. Hemi-BMP was detected as described above. Unit resolution was used in both MS1 and MS2, the collision energy was set to 25 V for BMP and hemi-BMP, 15V for PG, and 28V for LPG.

Data processing was performed with Agilent MassHunter quantitative analysis software version 10.1 and Agilent MassHunter qualitative analysis software version 10.0. Data were normalized for recovery, extraction and ionization efficacy by calculating analyte/internal standard ratios (AU); data are expressed as AU/µg protein or quantified via external calibration using BMP 36:2, hemi-BMP 54:3, PG 36:2, LPG 18:1 (Avanti Polar Lipids); and expressed as mol/g tissue or mol/g protein.

### Fluorescence microscopy

For microscopy, 3 x 10^5^ COS-7 cells were seeded in 35mm glass bottom dishes (Ibidi, Cat#81218-200). After 24h, cells were treated with 50 µM of indicated phospholipids and incubated overnight. Subsequently, cells were washed with PBS and incubated with fresh medium containing 1 µM LysoSensor™ Green DND-189 (Thermo Fisher Scientific, Cat#L7535) for 1h. Then, cells were washed with PBS and fresh medium was added containing 1 µg/ml Hoechst 33342 (Abcam, Cat# ab228551) for 10 min. Cells were washed with PBS and fresh medium was added.

Fluorescence microscopy was performed using a Leica SP8 confocal microscope with spectral detection (Leica Microsystems, Inc.), and an HC PL APO CS 63x/1.2 NA water immersion objective. LysoSensor^TM^ was excited at 488 nm, and emission was detected between 500-530 nm. Hoechst was excited at 350 nm, and emission was detected between 460 nm. Consistent imaging settings were applied for all samples.

The images were processed using the open-source software Fiji^38^ and the Leica LAS X software. Therefore, channels were split and a maximum intensity projection was generated for each channel. The background was subtracted using a rolling ball radius of 100 pixels for Hoechst and 50 pixels for LysoSensor^TM^ structures. The “Gaussian Blur” filter was applied on the Hoechst channel to enhance nuclei detection. Threshold values were set individually for each channel and kept constant for all images throughout all conditions. Based on the threshold, a mask was created and the “Watershed” algorithm was applied before particles were analyzed using the in-built “Analyze particle” function, with a particle size ranging from 5 to infinity for Hoechst and 0 to infinity for LysoSensor^TM^ structures. To determine the average LysoSensor^TM^ area/per cell, positive areas were summed and divided by the number of nuclei in each image.

### Quantification and statistical analysis

Data are presented as mean ± standard deviation (SD) unless stated otherwise. Differences in means were considered statistically significant for *p* < 0.05 (*), *p* < 0.01 (**), and *p* < 0.001 (***). The exact value and representation of n can either be found in the method section or in the figure legends. Statistical details can be found in the figure legends. Statistical analysis was performed using Prism version 9.5.1 (GraphPad).

## Results

### Accumulation of hemi-BMP and LPG precedes BMP synthesis

It is well established that the supplementation of mammalian cells with PG causes BMP accumulation and promotes the synthesis of intralumenal membranes in LE/lysosomes ^31,39,40^. Consistent with these studies, we observed that di-oleoyl PG [1,2-dioleoyl-sn-glycero-3-phospho-(1’-rac-glycerol), PG] increases BMP levels in various cell lines (**Figure 1A**). PG promoted the formation of acidic organelles in COS7 cells, which was not observed with di-oleoyl phosphatidylcholine (PC) and phosphatidylethanolamine (PE) (**Figure S1A&B**).

**Figure 1.**
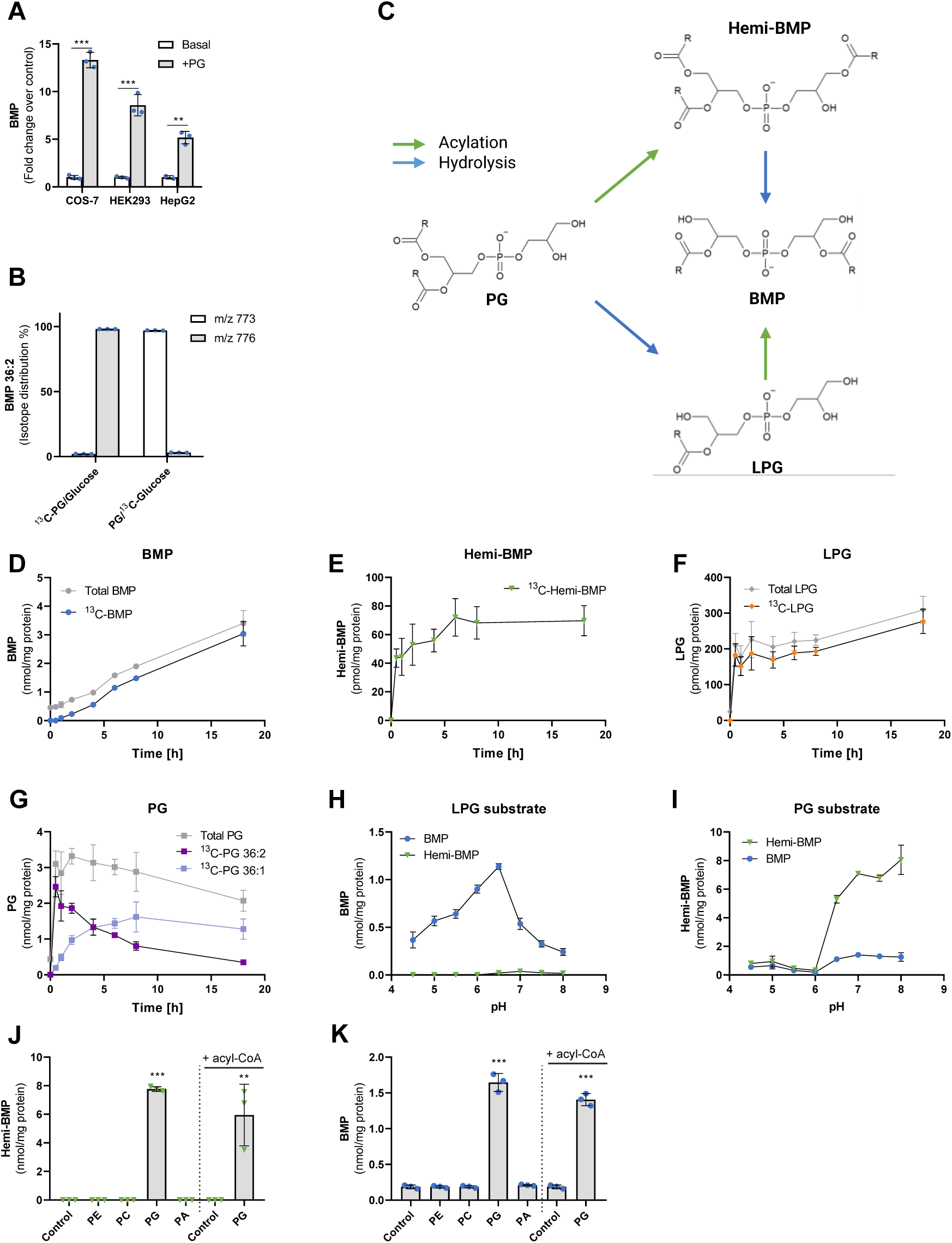
Accumulation of hemi-BMP and LPG precedes BMP synthesis. (**A**) Di-oleoyl PG supplementation (50 µM) induces BMP formation in the indicated cell lines (n=3). (**B**) Incorporation of head-group labeled di-oleoyl ^13^C-PG and UL-^13^C-glucose into BMP stores of COS-7 cells. M/z=773 and m/z=776 correspond to unlabeled and labeled di-oleoyl BMP, respectively (n=3). (**C**) Schematic depiction of BMP synthesis starting from its precursor PG. Acyl group positions and the stereoconfiguration of metabolites are described in the discussion section. (**D-G**) Time course of cellular BMP, hemi-BMP, LPG, and PG concentrations in HEK293 cells upon ^13^C-PG supplementation (n=4). Total (labeled plus unlabeled) as well as ^13^C-labeled metabolites were determined by LC-MS. (**H, I**) pH-dependent formation of BMP and hemi-BMP using Expi293F cell lysates as source of enzymatic activity and 1-oleoyl-LPG or di-oleoyl-PG as substrate (n=3). (**J, K**) Hemi-BMP and BMP formation by Expi293F lysates using the indicated substrates (n=3). All phospholipids were esterified with oleic acid. PG-induced formation of hemi-BMP/BMP was detected in the absence and presence of oleoyl-CoA (100 µM). Data are presented as mean ± SD. Statistically significant differences were determined by multiple unpaired *t*-tests for (A) and one-way ANOVA for (J) and (K), corrected for multiple comparisons against control by Bonferroni post hoc test (levels of statistically significant differences are: \*\**p* < 0.01, and \*\*\**p* < 0.001).

To study the conversion of PG to BMP in more detail, we monitored the incorporation of ^13^C-labeled di-oleoyl PG (^13^C-PG), containing uniformly-labeled (UL)-^13^C-glycerol as head group, into cellular BMP stores of COS-7 cells. Mass spectrometric analysis revealed that 98% of cellular di-oleoyl BMP exhibited a shift in the mass-to-charge ratio (m/z) from 773 to 776, demonstrating that the ^13^C-labeled head group of PG was efficiently incorporated into BMP. To investigate the exchange of glycerol moieties of newly formed BMP, we incubated cells with unlabeled PG and UL-^13^C-glucose, which delivers glycerol-3-phosphate for *de novo* lipid synthesis. After 18h of incubation, ∼ 2% of BMP contained the label, suggesting that the conversion of PG into BMP was not accompanied by a substantial exchange of backbone glycerol moieties (**Figure 1B**). This observation indicates that BMP synthesis does not require the degradation and re-synthesis of the di(glycero)phosphate backbone.

BMP synthesis necessitates the acylation of the head-group of PG or lyso-PG (LPG). PG acylation may lead to the formation of the intermediate acyl-PG (designated as hemi-BMP), which can be converted into BMP via a hydrolytic step. Alternatively, PG is first hydrolyzed to LPG and subsequently acylated (**Figure 1C**). To study BMP precursor formation in a cellular system, HEK293 cells were supplemented with ^13^C-PG. Subsequently, we monitored total and ^13^C-labeled BMP, hemi-BMP, LPG, and PG content in a time-course experiment. The ^13^C-label was efficiently incorporated into all newly formed metabolites (**Figure 1D-G**). The cellular BMP content increased in a linear manner during incubation (**Figure 1D**). PG, hemi-BMP and LPG accumulated already 30 min after PG supplementation and subsequently remained at elevated levels (**Figure 1E-G**). Thus, hemi-BMP and LPG accumulation precedes BMP synthesis, indicating that both metabolites can act as BMP precursors. PG 36:2 was converted to PG 36:1 during prolonged incubation (**Figure 1G**), which was not observed for BMP (**Figure S1C**).

For the initial biochemical characterization of BMP-synthesizing enzymes in human cells, we used lysates of Expi293F cells (derived from the HEK293 cell line) as source of enzymatic activity and different phospholipids as substrates. All lipid substrates were esterified with oleic acid. These experiments revealed that cell lysates contain enzymatic activities converting LPG into BMP and PG into hemi-BMP. LPG was most efficiently acylated in the acidic pH range with an optimum of 6.5, and the reaction did not generate hemi-BMP (**Figure 1H**). The pH dependence of BMP formation reflected exactly that of the recently identified lysosomal BMP synthase CLN5, which uses LPG as both donor and acceptor^22^. In comparison, highest hemi-BMP synthase activity was observed in the neutral and slightly alkaline pH range (**Figure 1I**). Hemi-BMP synthase activity was strongly reduced at pH values ≤ 6.0, indicating that PG acylation occurs outside of acidic organelles. The reaction also generated low amounts of BMP, which may derive from the degradation of hemi-BMP by cellular hydrolases. Hemi-BMP and BMP formation was not observed in the presence of phospholipid species other than PG (**Figure 1J, K**). Importantly, enzyme activity was not stimulated by the addition of acyl-CoA (**Figure 1J, K**), indicating that hemi-BMP can be synthesized by transacylation reactions transferring FAs from donor lipids to PG. Taken together, our observations suggest that the conversion of PG to BMP requires transacylation and hydrolase reactions, with either hemi-BMP or LPG as intermediate.

### Members of the cytosolic phospholipase A2 group IV enzyme family catalyze hemi-BMP and BMP synthesis

To investigate the relevance of the PG - hemi-BMP - BMP pathway (**Figure 1C**), we first attempted to find enzymes that catalyze hemi-BMP synthesis. Lipid transacylase reactions are catalyzed by enzymes of different protein families including the α/β hydrolase superfamily^41^, the patatin-like phospholipase domain-containing (PNPLA) family^42^, and the H-RAS-like suppressor family^43^. To identify hemi-BMP synthases, we screened a library of 206 mouse enzymes comprising most known lipid hydrolases and transacylases encoded by the mouse genome as well as structurally related proteins with unknown function. Proteins were expressed in Expi293F cells and cell lysates were used to screen for hemi-BMP synthase activity. The expression of 91% of the His- or Strep-tagged recombinant proteins was confirmed either by Western blotting analysis or using activity-based assays (Supplemental Table S2, DOI: 10.17632/hcs7crxk3z.1). Strikingly, the top hits of the hemi-BMP synthase screen were assigned to two members of the cytosolic phospholipase A2 group IV protein family (PLA2G4D and E) (**Figure 2A**).

**Figure 2.**
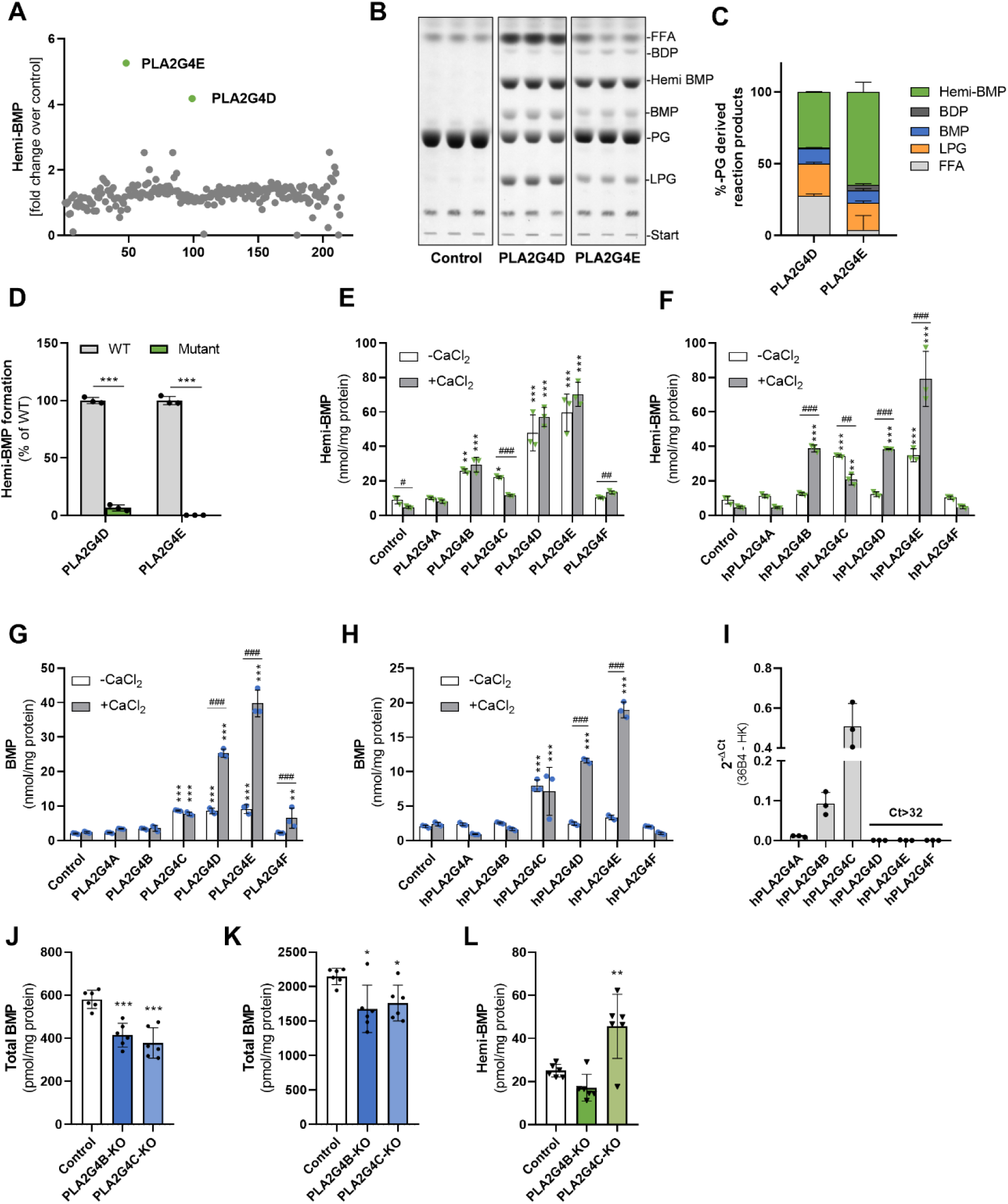
Members of the cytosolic phospholipase A2 group IV enzyme family catalyze hemi-BMP and BMP synthesis. (**A**) Screening for hemi-BMP synthase activity using a lipid-hydrolase/transacylase enzyme library (n=1). (**B**) TLC analysis of the reaction products of semi-purified PLA2G4D and PLA2G4E incubated with di-oleoyl PG. (**C**) Densitometric quantitation of (B) (n=3). (**D**) Hemi-BMP synthase activity of wild-type (WT) and mutant enzymes. The active serine of PLA2G4D and PLA2G4E was replaced with an alanine (S370A and S420A, respectively). Lysates of Expi293F cells expressing WT and mutant variants were used as source of enzymatic activity (n=3). (**E, F**) Hemi-BMP synthase activity of mouse and human PLA2G4 (hPLA2G4A-F) orthologues (**G, H**) BMP synthase activity of mouse and human PLA2G4 orthologues. In (E)-(H), lysates of Expi293F cells expressing recombinant enzymes were used as source of enzymatic activity. Di-oleoyl PG was used as substrate and assays were performed in the absence and presence of 2 mM CaCl_2_(n=3, representative of three independent experiments). (**I**) Gene expression analysis of PLA2G4 family members in HEK293 cells (n=3). (**J**) Basal BMP content of control, *PLA2G4B*-KO, and *PLA2G4C*-KO cells (n=6, representative of three independent experiments). (**K, L**) BMP and hemi-BMP content of control, *PLA2G4B*-KO, and *PLA2G4C*-KO cells after incubation with PG (50 µM) for 12h (n=6, representative of three independent experiments). Data are shown as mean ± SD. Statistically significant differences were determined by multiple unpaired *t*-tests for (D), two-way ANOVA for (E-H), and one-way ANOVA for (J-L), followed by corrections for multiple comparisons (* = compared to the respective control, ^#^ = comparison between groups) using Bonferroni post hoc test (levels of statistically significant differences are: *^,#^*p* < 0.05, **^,##^*p* < 0.01, and ***^,###^*p* < 0.001).

To characterize the activities of PLA2G4D and -E in more detail, we enriched recombinant His-tagged mouse enzymes by affinity chromatography (**Figure S2A**), incubated the partially purified proteins with PG, and analyzed the reaction products by TLC. Both enzymes generated hemi-BMP, BMP, LPG, FAs, and also low amounts of bis(diacylglycero)phosphate (BDP), representing fully acylated PG (**Figure 2B**). Semi-quantitative analysis of the reaction products revealed that hemi-BMP is the major acylation product (**Figure 2C**). In comparison to PLA2G4D, PLA2G4E released lower amounts of FAs suggesting a preference for transacylation over hydrolytic reactions under the employed experimental conditions. These experiments confirm that PLA2G4D and E possess both PG hydrolase and transacylase activity. PG can act as an acyl donor and acceptor in the transacylation reaction. The replacement of the active serine with an alanine abolished hemi-BMP synthase activity of both enzymes (**Figure 2D**).

The PLA2G4 family comprises 6 members that share structural similarities (PLA2G4A - F, commonly also referred to as cPLA2α - ζ)^44^. These proteins are characterized by a catalytic domain that is similar to the patatin-like domain of the plant lipid hydrolase patatin. All family members, except PLA2G4C, contain an N-terminal Ca^2+^-binding C2 domain involved in the regulation of enzyme activity^44^ (**Figure S2B**). We thus compared the activity of all members of this family in the absence and presence of CaCl_2_. Assays were performed using Expi293F lysates expressing the recombinant mouse and human proteins and PG as substrate. Protein expression was verified by Western blotting analysis (**Figure S2C&D**). Using mouse enzymes, we observed increased hemi-BMP formation in lysates containing PLA2G4B-F. CaCl_2_ barely stimulated hemi-BMP formation, while causing reduced activity of PLA2G4C (**Figure 2E**). Human PLA2G4 enzymes exhibited similar activities. In contrast to mouse enzymes, addition of CaCl_2_ promoted the hemi-BMP synthase activity of PLA2G4B, -D, and -E, while PLA2G4C activity was again decreased (**Figure 2F**). PLA2G4 mediated reactions also resulted in increased BMP formation, whereby CaCl_2_ strongly promoted PLA2G4D and -E activity of mouse and human orthologues (**Figure 2G&H**).

The transacylase activities of mouse PLAG4B-F were confirmed using partially purified protein fractions and TLC-based lipid analysis in the presence and absence of CaCl_2_ (**Figure S2E-J**). Using PG as substrate, all enzyme-enriched fractions generated hemi-BMP, of which PLA2G4F was activated by CaCl_2_ (**Figure S2E&F**). PLA2G4D, E, & F also generated BMP, likely through the hydrolysis of hemi-BMP (**Figure S2E&G**). In addition, all purified enzymes except PLA2G4F were capable of acylating 1-oleoyl LPG leading to the formation of BMP and hemi-BMP, albeit at a lower rate in comparison to PG acylation (**Figure S2H-J**). An exception is PLA2G4C, which showed higher activity using LPG as substrate.

To compare the hydrolytic activities of mouse PLA2G4 enzymes, we incubated cell lysates containing recombinant proteins with tri-oleoyl hemi-BMP and di-oleoyl BMP as substrates, and monitored the release of FAs. PLA2G4A-D showed hemi-BMP hydrolase activity (**Figure S2K**), while specifically PLA2G4B and -D degraded BMP (**Figure S2L**). Addition of CaCl_2_ barely affected hemi-BMP hydrolysis, but increased BMP degradation by PLA2G4B and -D (**Figure S2K&L**). The hemi-BMP hydrolase/transacylase activities of PLA2G4B-F were confirmed using partially purified protein fractions and TLC analysis. Depending on the substrate specificity of different enzymes, digestion of hemi-BMP resulted in the formation of BMP, PG, LPG, and BDP (**Figure S2M-Q**).

Taken together, our observations suggest that specific members of the PLA2G4 family possess both transacylase and hydrolase activity leading to the formation or degradation of hemi-BMP/BMP. To study the role of endogenously expressed PLA2G4 enzymes in HEK293 cells, we first analyzed gene expression by qPCR analysis, which revealed that HEK293 cells primarily express PLA2G4B and C (**Figure 2I**). Accordingly, we investigated BMP metabolism in HEK293 cells lacking either of these enzymes. The CRISPR/Cas9 system was utilized to generate knockout cell lines through the introduction of a frame shift within the targeted exon, resulting in a premature stop-codon (**Figure S2R**). Under basal conditions, deletion of PLA2G4B or PLA2G4C decreased the cellular BMP content by 25% and 30%, respectively (**Figure 2J**). PG-induced BMP synthesis in both mutant cell lines was reduced by ∼ 20% (**Figure 2K**), due to a reduction of the most abundant BMP subspecies (**Figure S2S&T**). Unexpectedly, however, we found unchanged hemi-BMP levels in PLA2G4B-deficient cells and even increased levels in PLA2G4C-deficient cells (**Figure 2L**). We were not able to generate a double knockout cell line, since combined deletion of both PLA2G4C and B, either by *PLA2G4C* deletion in PLA2G4B-deficient cells or *vice versa*, resulted in cell death after transfection.

### Members of the pancreatic lipase family catalyze extracellular hemi-BMP and BMP synthesis

Our observation in mutant HEK293 cells strongly indicated that hemi-BMP can be synthesized independently of PLA2G4-catalyzed reactions. In further experiments, we observed that fetal bovine serum (FBS) contains substantial hemi-BMP synthase activity. This activity was abolished by heat inactivation or by the addition of the non-specific serine hydrolase inhibitor Orlistat^45^ (**Figure 3A**), suggesting that hemi-BMP is synthesized by serum enzymes. To identify the responsible enzymes, we monitored hemi-BMP formation in supernatants of Expi293F cells expressing transacylases/hydrolases of different enzymes families. This screen identified one enzyme that increased hemi-BMP synthesis 40-fold compared to the empty vector control (**Figure 3B**) and several other enzymes with lower activity. The top hit was annotated to endothelial lipase (gene name *Lipg*), a well-characterized serum phospholipase of the pancreatic lipase family^46^. The hemi-BMP synthase activity of an inactive LIPG variant (S169A) was strongly reduced in comparison to the wild-type control (**Insert Figure 3B**).

**Figure 3.**
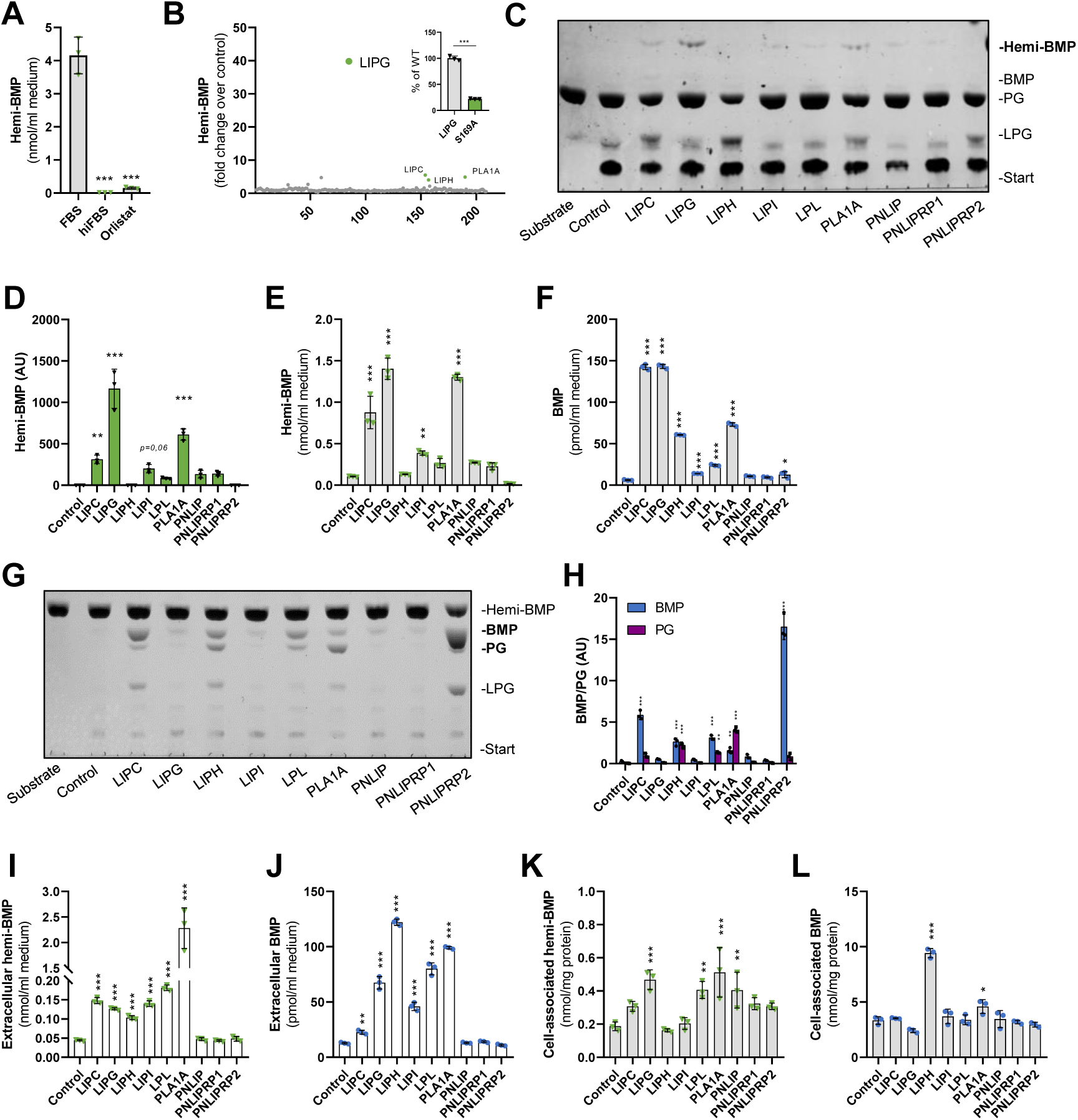
Members of the pancreatic lipase family catalyze extracellular hemi-BMP and BMP synthesis. (**A**) Hemi-BMP formation in DMEM/10% FBS, DMEM/10% heat-inactivated FBS (hiFBS), and DMEM/10%FBS/40 µM Orlistat (n=3) using di-oleoyl PG as substrate. (**B**) Screening for hemi-BMP synthase activity using supernatants of Expi293F cells expressing mouse lipid hydrolases/transacylases (n=1). The insert shows hemi-BMP synthase activity detected in supernatants of cells expressing wild-type LIPG and its catalytically inactive S169A variant (n=3). (**C**) Representative TLC showing hemi-BMP synthase activity of PNLIP family members in the presence of di-oleoyl PG. Conditioned (serum-free) media of Expi293F cells expressing recombinant enzymes were used as source of enzymatic activity. Note that BMP signals were barely visible. **(D**) Densitometric analysis of hemi-BMP formation shown in (C) (n=3). (**E, F**) Hemi-BMP and BMP synthase activity of PNLIP family members was confirmed by LC-MS-based analysis of reaction products (n=3). (**G**) Representative TLC showing hemi-BMP hydrolase activity of PNLIP family members. Conditioned (serum-free) media of Expi293F cells expressing recombinant enzymes were used as source of enzymatic activity. (**H**) Densitometric analysis of BMP and PG formation shown in (G) (n=3). (**I**, **J**) Extracellular and (**K**, **L**) cell-associated hemi-BMP and BMP content of HEK293 cells overexpressing indicated members of the pancreatic lipase family after 4h of PG supplementation. Experiments shown in (I-L) were performed in DMEM medium containing 10% heat-inactivated FBS (n=3) and are representative of two independent experiments. Data are presented as mean ± SD. Statistically significant differences were determined by one-way ANOVA, followed by corrections for multiple comparisons versus control using Bonferroni post hoc test (levels of statistically significant differences are: \**p* < 0.05, \*\**p* < 0.01, and \*\*\**p* < 0.001).

LIPG belongs to a group of enzymes, which share structural similarity with pancreatic lipase (PNLIP). Screening experiments indicated that several other enzymes of this family increase hemi-BMP formation (**Figure 3B**), although to a lower extent than LIPG. The mouse pancreatic lipase gene family comprises 9 proteins characterized by an N-terminal α/β hydrolase fold, which contains the catalytic site, and C-terminal domains mediating the interaction with lipid substrates and co-activators^47,48^ (**Figure S3A**). PNLIP and Pnlip-related protein 1 (PNLIPRP1) are primarily expressed in exocrine pancreatic cells, while PNLIPRP2 exhibits a broader expression spectrum and is also active in the circulation together with LIPG, hepatic lipase (LIPC), lipase H (LIPH), lipase I (LIPI), lipoprotein lipase (LPL), and phospholipase A1 member A (PLA1A).

To compare the activities of PNLIP enzymes, we monitored hemi-BMP and BMP formation by TLC in conditioned media of Expi293F cells secreting recombinant mouse enzymes. In addition to LIPG, we detected hemi-BMP synthase activity for several other enzymes of the PNLIP family (**Figure 3C&D**). These findings were verified in independent experiments by MS analysis confirming the highest hemi-BMP synthase activity for LIPG, followed by PLA1A, LIPC, and LIPI, respectively (**Figure 3E**). All pancreatic lipase family members, except PNLIP and PNLIPRP1, generated BMP at a lower extent than hemi-BMP (**Figure 3F**). To investigate whether these enzymes hydrolyze hemi-BMP to yield BMP, conditioned media were incubated with hemi-BMP as substrate and the reaction products were separated by TLC (**Figure 3G&H**). Indeed, several enzymes hydrolyzed hemi-BMP and generated both PG and BMP, suggesting that FAs are mobilized from both double and single acylated glycerol moieties of hemi-BMP. LIPC, LPL, and PNLIPRP2 preferentially generated BMP, while LIPH produced equimolar amounts of PG and BMP, and PLA1A mostly generated PG (**Figure 3H**). For other enzymes, hemi-BMP hydrolase activity was not or barely detectable. As LIPG did not exhibit hemi-BMP hydrolase activity, the observed BMP formation cannot derive from the hydrolysis of hemi-BMP. Therefore, we tested whether LPG can also act as acyl acceptor and acyl donor for LIPG. As shown in **Figure S3B&C**, LIPG was the only enzyme capable of acylating LPG, which led to the formation of both BMP and PG.

Next, we overexpressed all nine PNLIP family members in HEK293 cells and monitored hemi-BMP and BMP formation in cell culture supernatants and cells upon PG supplementation. Overexpression of LIPC, LIPG, LIPH, LIPI, LPL, and PLA1A increased extracellular hemi-BMP and BMP formation. Conversely, PNLIP, PNLIPRP1, and PNLIPRP2 had no effect (**Figure 3I&J**). After the 4 h incubation period, cell-associated hemi-BMP levels were significantly increased in LIPG-, LPL-, PLA1A-, and PNLIP-expressing cells (**Figure 3K**). LIPH- and PLA1A-expressing cells exhibited elevated cellular BMP levels (**Figure 3L**).

### PLA2G4 and PNLIP family members increase BMP formation in *CLN5*-deficient cells

Our observations suggest that cytosolic and secreted enzymes catalyze head group acylation of PG/LPG. However, this reaction can also occur in lysosomes^22^ indicating a redundant system. To discriminate between intra- and extra-lysosomal pathways, we investigated whether PLA2G4B-F and the most active secreted PNLIP enzymes can increase BMP synthesis in the absence of the lysosomal BMP synthase CLN5. HEK293 cells lacking CLN5 (*CLN5*-KO) were generated using the CRISPR/Cas9 system and *CLN5* deletion was confirmed by Sanger sequencing (**Figure S4A**) and Western blotting (**Figure 4A**). Consistent with previous observations, we found that *CLN5*-KO cells accumulate LPG (**Figure 4B**). Unexpectedly, however, BMP levels were unchanged (**Figure 4C**). Upon PG supplementation, LPG levels of *CLN5*-KO cells were increased by ∼ 3-fold compared to wild-type controls (**Figure 4D**). Conversely, BMP levels were reduced by 50% (**Figure 4E**), confirming that *CLN5*-KO cells have a reduced capacity to produce BMP. Overexpression of PLA2G4 members did not affect BMP accumulation in *CLN5*-KO cells under basal conditions, while LIPC and PLA1A led to a moderate increase (**Figure 4F**). In the presence of PG, PLA2G4D and all tested PNLIP members increased BMP synthesis in *CLN5-*KO cells to wild-type levels (**Figure 4G**). Thus, cytosolic and secreted transacylases are capable of promoting BMP synthesis independently of CLN5.

**Figure 4.**
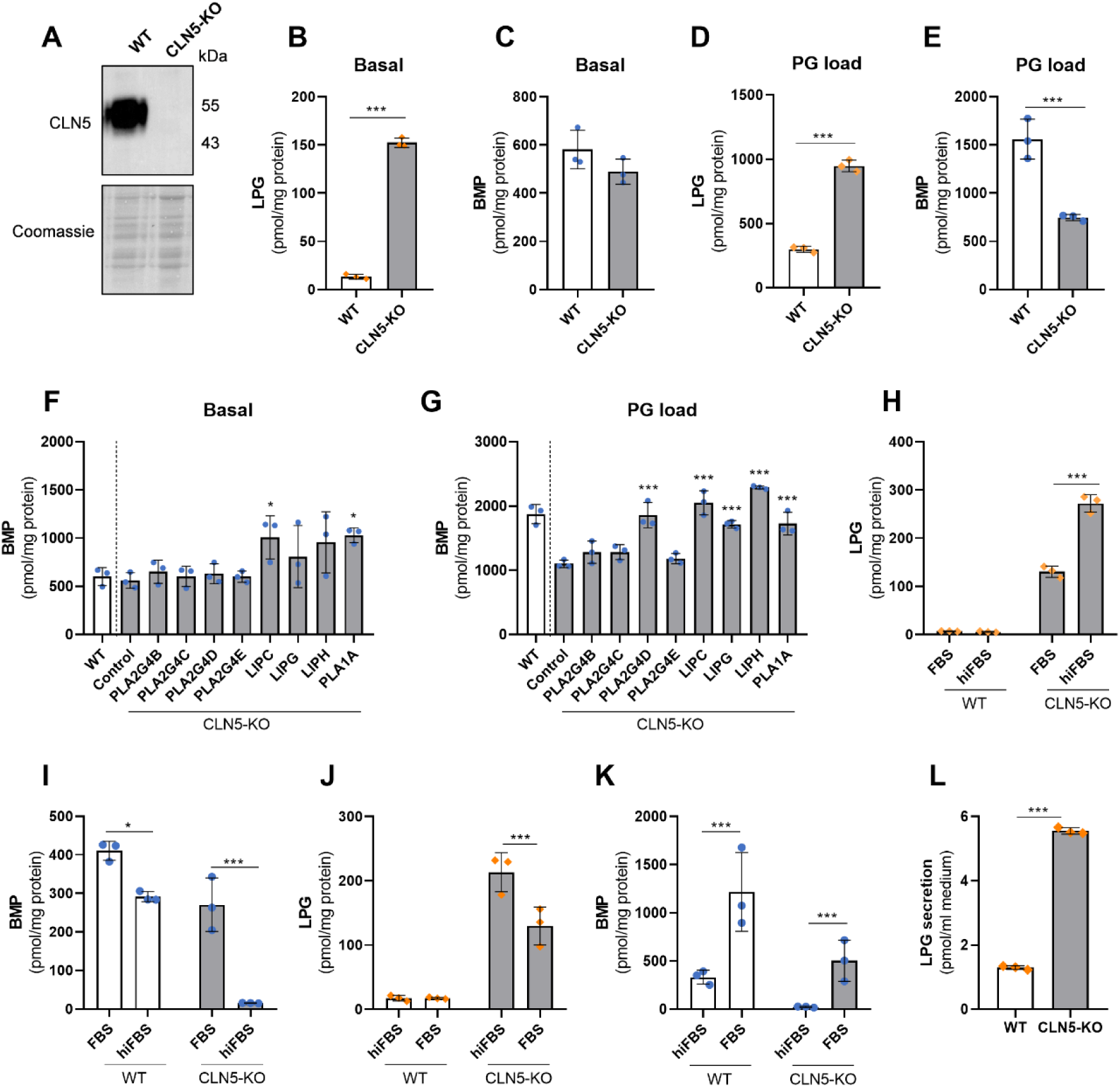
Members of the PLA2G4 and PNLIP family promote BMP synthesis independently of CLN5. (**A**) Verification of *CLN5* knockout in HEK293 cells by Western blot analysis. The Coomassie-stained blot was used as loading control. (**B, C**) Basal LPG and BMP content in WT and *CLN5*-KO cells (n=3). (**D, E**) BMP content of WT and *CLN5*-KO cells after supplementation with di-oleoyl PG (50 µM) for 6h (n=3). (**F, G**) BMP content of CLN5-KO cells upon overexpression of indicated PLA2G4 and PNLIP family members under basal conditions and after 8h PG supplementation (50 µM, n=3). The WT control was transfected with empty vector. Note that cells used for experiments B-G were cultivated in DMEM containing 10% FBS. For experiments (6-8 h), FBS was replaced by hiFBS to avoid serum-mediated acylation reactions. (**H, I**) LPG and BMP of WT and CLN5-KO cells cultivated for two weeks in DMEM medium containing either 10% FBS or hiFBS (n=3). (**J, K**) LPG and BMP content of WT and *CLN5*-KO cells cultured in DMEM with 10% hiFBS for one week and subsequently in DMEM with 10% FBS for another week (n=3). (**L**) Secretion of LPG by WT and *CLN5*-KO HEK293 cells (n=3). Cells were incubated in DMEM containing 2% BSA for 24 h and lipids were extracted from lyophilized conditioned media. Data are presented as mean ± SD and representative for at least two independent experiments. Statistically significant differences were determined by unpaired Student’s *t*-test (B-E, L), one-way ANOVA (F, G), and two-way ANOVA (H-K) followed by corrections for multiple comparisons versus control using Bonferroni post hoc test (levels of statistically significant differences are: \**p* < 0.05, \*\**p* < 0.01, and \*\*\**p* < 0.001).

Recent data suggested that *CLN5*-KO cells exhibit severe BMP deficiency^22^. To explain this discrepancy with our measurements, cells were cultured for 14 days in medium containing either FBS or heat-inactivated FBS (hiFBS) lacking transacylase activity. Using hiFBS, the LPG content remained low in wild-type (WT) cells and further increased ∼ 2-fold in *CLN5*-KO cells in comparison to cells cultured in FBS (**Figure 4H**). The BMP content of WT cells was reduced by 25% in the presence of hiFBS, while *CLN5*-KO cells almost completely lost their BMP stores under these conditions (**Figure 4I**). A switch from hiFBS- to FBS-containing culture medium had opposite effects. LPG levels decreased in *CLN5*-KO cells (**Figure 4J**), while BMP increased in both WT and *CLN5*-KO cells (**Figure 4K**). These observations indicate that cells secrete PG or LPG, which is acylated by serum transacylases and reabsorbed as hemi-BMP or BMP. Analysis of conditioned media revealed that cells secrete LPG, while PG was not detectable. LPG secretion was increased by ∼ 5-fold in *CLN5*-KO cells compared to WT controls (**Figure 4L**). Together, our data suggest that BMP synthesis in *CLN5*-KO cells strongly depends on extracellular generation of BMP precursors by serum transacylases. This process also enhances BMP synthesis in WT cells.

### Serum transacylases produce hemi-BMP and BMP *in vitro* and *in vivo*

Several members of the pancreatic lipase family possess transacylase activity generating hemi-BMP or BMP, hydrolase activity converting hemi-BMP into BMP, or both activities (**Figure 3**). Under physiological conditions, these enzymes are active in the circulation and might therefore cooperatively generate BMP or precursors thereof. To investigate hemi-BMP/BMP synthesis in by serum transacylases, we incubated plasma of heparinized mice with LPG or PG and monitored the formation of acylated products. Since LIPG was identified as the most potent transacylase, we also compared hemi-BMP/BMP synthesis in plasma of wild-type and *Lipg-*deficient mice (*Lipg*-ko). Incubation of mouse plasma with both LPG and PG led to the formation of BMP (**Figure 5A**). We could not detect differences in BMP formation between genotypes. However, BMP generated in *Lipg-*deficient plasma showed an altered FA composition (**Figure S5A&B**). In the presence of LPG, we also observed PG formation indicating that the substrate can be acylated on both glycerol moieties (**Figure 5B**). Both lipid substrates led to the synthesis of hemi-BMP, whereby hemi-BMP formation was ∼ 10-fold higher in the presence of PG in comparison to LPG (**Figure 5C**). Hemi-BMP synthesis in *Lipg* -deficient plasma was reduced by ∼ 40% in the presence of LPG and by 30% using PG as substrate (**Figure 5C**). Notably, LPG- and PG-derived hemi-BMP strongly differed in FA composition (**Figure S5C&D**). The reduced synthesis of hemi-BMP levels by *Lipg* -deficient plasma was mainly due to a decrease of the major hemi-BMP species 52:2 and 54:3 detected in the presence of LPG and PG, respectively (**Figure S5C&D**). Thus, serum transacylases can mediate head group acylation of both LPG and PG. Their conversion into hemi-BMP/BMP is catalyzed by LIPG and other transacylases.

**Figure 5.**
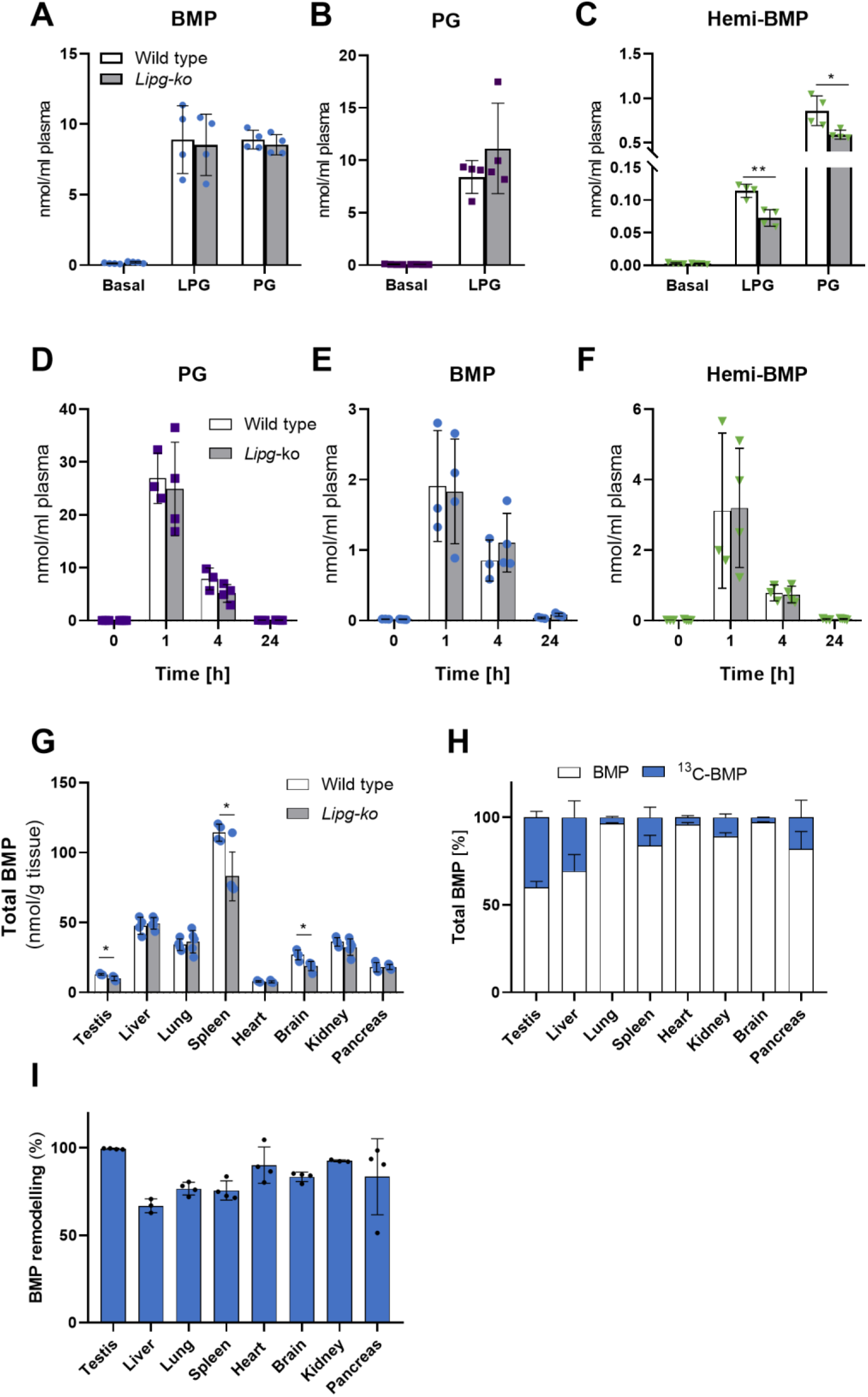
Serum transacylases produce hemi-BMP and BMP *in vitro* and *in vivo*. (**A-C**) BMP, PG, and hemi-BMP synthesis by heparin-plasma of wild-type and *Lipg*-ko mice using *sn-*1-oleoyl LPG or di-oleoyl PG as substrate (n=4). (**D-F**) Plasma PG, BMP, and hemi-BMP content of mice after intraperitoneal injection of di-oleoyl PG (wild-type n=3; *Lipg*-*k*o n=4) (5 mg/mouse). Blood samples of wild-type and *Lipg*-ko mice were collected before injection and at the indicated time points after injection (n=3). (**G**) Total tissue BMP content of wild-type (n=4) and *Lipg*-ko mice (n=5). Data are presented as mean ± SD. (**H**) Incorporation of ^13^C-labeled di-oleoyl PG into tissue BMP stores. (**I**) FA remodeling of ^13^C-labeled di-oleoyl BMP (n=3-4). In (H) and (I), tissues were collected 4h after intraperitoneal injection of 0.2 mg ^13^C-PG/mouse (n=3-4). Statistically significant differences were determined by unpaired two-tailed Student’s *t*-test, followed by corrections for multiple comparisons using the Holm-Sidak post hoc test (levels of statistically significant differences are: \**p* < 0.05, \*\**p* < 0.01, \*\*\**p* < 0.001).

To monitor PG conversion *in vivo*, we treated mice with di-oleoyl PG and measured plasma hemi-BMP and BMP levels. Basal PG levels were barely detectable and PG treatment led to a strong increase in circulating PG after 1 h (**Figure 5D**). Within the next 3 h, plasma PG decreased by ∼70% due to degradation, cellular uptake, or conversion into other lipids, and after 24 h, PG concentrations returned to basal levels. Elevated PG levels were accompanied by an increase in circulating BMP (**Figure 5E**) and hemi-BMP levels (**Figure 5F**). BMP and hemi-BMP content together accounted for ∼ 20% of PG, suggesting that a substantial portion of PG is converted into these metabolites. We did not observe an attenuated conversion of circulating PG into hemi-BMP/BMP in *Lipg*-ko mice in comparison to wild-type controls (**Figure 5D-F**), implicating that other serum transacylases largely compensate for the lack of LIPG activity. Notably, however, we observed that independent of the genotype more than ∼ 60% of newly formed hemi-BMP/BMP were acylated with palmitic acid (16:0) or stearic acid (18:0), indicating that FAs derive from the *sn*-1 position of donor phospholipids (**Figure S5E-F**), which are typically esterified with saturated FAs.

Next, we compared tissue BMP levels of wild-type and *Lipg-*ko mice. Despite the overlapping functions of intra- and extracellular transacylases, we found moderately reduced BMP levels in spleen, testis, and brain of *Lipg*-ko mice (**Figure 5G**). Tissue distribution and tissue-specific changes of BMP subspecies are shown in **Figure S5G-N**.

Finally, to investigate BMP synthesis *in vivo*, we treated wild-type mice with ^13^C-PG and monitored the incorporation of the label into the BMP moiety of different tissues after 4 h (**Figure 5H**). The head group of PG was efficiently incorporated into BMP of testis, where the labeled fraction accounted for 40% of total BMP, followed by liver (20%), pancreas (16%), spleen (12%), and kidney (8%), while minor incorporation was observed in lung, heart, and brain (< 3%). To monitor FA remodeling of BMP, we calculated the percentage of ^13^C-labeled BMP subspecies, where at least one of the original oleic acids of ^13^C-PG was replaced by other FAs. Remarkably, FA remodeling reached almost 100% in testis and was also very effective in other tissues (**Figure 5I**). Taken together, our data suggest that circulating PG is efficiently incorporated into the BMP pool of different tissues, followed by tissue-specific FA remodeling.

## Discussion

BMP was first described by Body and Gray in 1967^49^. Since its discovery, the molecular pathway of BMP synthesis remained incompletely understood. Here, we show that extra- and intracellular pathways promote BMP synthesis.

The synthesis of BMP necessitates the acylation of the head group of the precursor lipids PG or LPG. The initial characterization of cellular transacylation activities revealed two significant insights into BMP synthesis: firstly, that LPG and PG are preferentially acylated in the acidic and neutral pH range, respectively, suggesting the existence of distinct synthesis pathways; and secondly, that cells exposed to PG accumulate hemi-BMP and LPG, indicating that both metabolites can serve as intermediates of BMP synthesis. These observations indicate that BMP synthesis can be initiated either by acylation of PG to hemi-BMP followed by hydrolysis to BMP, or by hydrolysis of PG to LPG followed by acylation to BMP.

Recent observations suggested that LPG is produced from PG in acidic organelles by lysosomal phospholipase A2 (PLA2G15)^50^ and subsequently acylated by the lysosomal transacylase CLN5^22^, resulting in BMP formation. CLN5 utilizes LPG as both acyl donor and acceptor, and does not produce hemi-BMP^22^. Therefore, it remained unclear how hemi-BMP is synthesized. We screened an enzyme library for PG transacylation activity and identified members of the PLA2G4 and PNLIP enzyme families as hemi-BMP synthases. All identified enzymes, except PLA2G4C, preferred PG over LPG as acyl acceptor, confirming that neutral transacylases preferentially produce hemi-BMP. Several of the identified intracellular and secreted enzymes also showed hydrolase activity converting hemi-BMP into BMP and BMP into LPG, indicating that they can mediate BMP synthesis and degradation. Within the PLA2G4 family, transacylase activities were previously reported for PLA2G4C and -E. PLA2G4C mediates the acylation of lysophospholipids^51^, while PLA2G4E acylates phosphatidylethanolamine (PE), resulting in the formation of N-acyl PE, the precursor of the endocannabinoid *N*-arachidonoyl ethanolamine (anandamide) and other N-acylated ethanolamines^52^. Interestingly, PLA2G4E associates with endocytic vesicles and is involved in the regulation of trafficking within endocytic and recycling routes^53^. Research on PNLIP enzymes has extensively focused on their catabolic functions, while transacylase reactions have received little attention. These enzymes possess central functions in lipid degradation in the intestinal lumen and in the circulation^47,54,55^. In accordance with our findings, PNLIPRP2 was previously shown to hydrolyze BMP and hemi-BMP^56^. Transacylase activity was reported for LIPC, which can mediate the acylation of dolichol^57^. Additionally, PNLIP is frequently used for the *in vitro* synthesis of lipids^58^.

Our screening experiments identified cytosolic and secreted enzymes as hemi-BMP/BMP synthases suggesting that extra-lysosomal transacylation reactions can deliver BMP precursors that are subsequently transferred to LE/lysosomes. To distinguish between acidic and neutral pathways, we monitored BMP synthesis in *CLN5*-KO cells. These cells show a reduced capacity to produce BMP upon PG supplementation. Notably, overexpression of PLA2G4D, LIPC, LIPG, LIPH, and PLA1A increased BMP levels to wild-type levels, suggesting that secreted, cytosolic, and lysosomal transacylases promote BMP synthesis (summarized in **Figure 6A**). Furthermore, we found that the cellular BMP levels are strongly influenced by cell culture conditions. *CLN5*-KO HEK293 cells exhibit normal BMP levels under standard conditions and lose BMP stores in the presence of hiFBS lacking transacylase activity. Our observations suggest that cells secrete LPG, which can be acylated by serum enzymes and subsequently reabsorbed. Apparently, this secretion-acylation-reuptake cycle can compensate for the lack of lysosomal transacylation activity and also promotes BMP synthesis in wild-type HEK293 cells.

**Figure 6.**
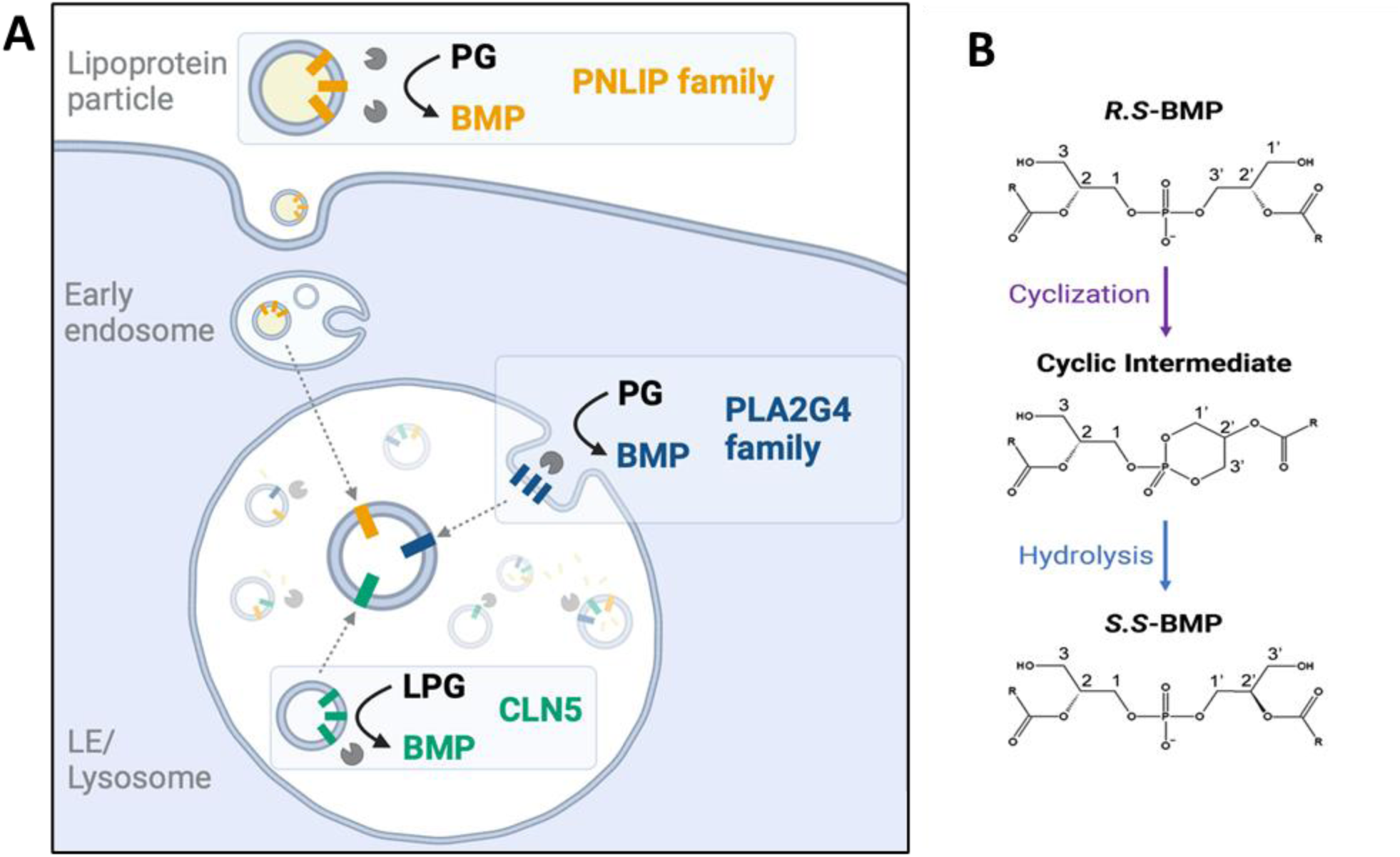
BMP synthesis is promoted by functionally overlapping pathways. (**A**) The acylation of the head group of the precursor lipids PG or LPG is a necessary step in BMP synthesis. This reaction can be catalyzed by secreted (PNLIP family), cytosolic (PLA2G4 family), and lysosomal (CLN5) enzymes. The acylation of natural *R,S* - PG/LPG delivers BMP precursors with *R,S* - configuration, which are then subjected to stereoconversion and FA remodeling. These reactions deliver mature *S,S* - BMP, which is primarily acylated in *sn*-2 positions. The mechanism and site of stereoconversion remain unknown. One possible mechanism is the formation of a cyclic intermediate followed by its stereospecific hydrolysis, as depicted in (**B**).

*CLN5* loss-of-function mutations are associated with the accumulation of autofluorescent lipopigments (lipofuscin) in lysosomes in the brain and peripheral tissues^59^. The phenotype in peripheral tissues is much milder than neurological symptoms, although CLN5 is highly expressed in many peripheral tissues^59^. The cytosolic and secreted enzymes identified in this study increase BMP synthesis independently of CLN5 and may therefore counteract BMP deficiency in CLN5-deficient tissues. In particular, access to circulating hemi-BMP and BMP, which may derive from dietary or endogenously produced LPG or PG, could attenuate the phenotype. PG is a highly abundant lipid in plant chloroplasts and bacterial membranes^60^ and it is reasonable to assume that this lipid is present in substantial amounts in the human diet. However, PG is found only at low concentrations in human serum^60^ indicating that it is rapidly degraded or converted into other lipids. Our *in vivo* experiments in mice revealed that a substantial portion of circulating PG is converted to hemi-BMP and BMP. Previous work demonstrated that BMP is present in lipoproteins^61,62^ and endosome-derived extracellular vesicles^63^, implicating that it can enter the endolysosomal pathway via receptor-mediated endocytosis. BMP is highly resistant to lysosomal hydrolases^64,65^ and may therefore accumulate in acidic organelles.

In mouse studies, we observed that about 50% of PG-derived circulating hemi-BMP and BMP species were esterified with palmitic or stearic acid. Considering that saturated FAs are typically present at the *sn*-1 position of phospholipids, this observation is consistent with the previously reported *sn*-1(3) regioselectivity of the PNLIP family^66^. Studies with stably labeled PG revealed that the glycerol head group of PG is efficiently incorporated into the BMP pool of different tissues. This may occur through the uptake of extracellularly formed hemi-BMP/BMP or through the direct uptake of PG/LPG, followed by intracellular conversion to BMP. In both cases, the uptake must be followed by tissue-specific remodeling of FA moieties, since the FA composition of ^13^C-labeled BMP extracted from tissues strongly differs from the applied di-oleoyl PG. We previously reported that mouse tissues exhibit very characteristic BMP profiles. For example, the predominant BMP species in muscle and adipose tissue depots are esterified with oleic (18:1) and linoleic acid (18:2), while brain and testis contain BMP almost exclusively esterified with docosahexaenoic acid (22:6)^67^. These differences in FA composition suggest that BMP synthesis and remodeling are catalyzed by intracellular transacylases/hydrolases that exhibit selectivity for specific FA species. PLA2G4 family members differ in their cell- and tissue-specific expression patterns^44^ and can catalyze BMP synthesis as well as degradation. It remains to be investigated whether these enzymes show FA preferences and thus determine BMP composition in different tissues.

LIPG was identified as the most potent extracellular PG transacylase. We found that this enzyme is responsible for ∼ 30% of the hemi-BMP synthase activity detected in heparinized mouse plasma and affects the FA composition of both hemi-BMP and BMP. *Lipg*-ko mice have moderately reduced BMP content in testis, brain, and spleen, suggesting that LIPG promotes BMP synthesis in a tissue-specific manner. These tissues have low CLN5 mRNA and protein levels compared to other tissues such as the liver, kidney, and heart^68,69^, and may therefore react more sensitive to *Lipg*-deficiency. However, the conversion of circulating PG into hemi-BMP/BMP was not affected in *Lipg*-ko mice, most likely due to the overlapping function of serum transacylases. Overall, our observations suggest that in addition to LIPG, other serum lipases/transacylases substantially contribute to hemi-BMP/BMP formation in the circulation.

BMP isolated from mammalian cells has an *sn*-1-glycerophosphate-*sn*-1′-glycerol (*S,S*) backbone stereoconfiguration, whereas PG shows *sn*-3:*sn*-1 (*R,S*) configuration^70^. Acyl chains of natural BMP are predominantly present on the 2- and 2’-positions of the backbone^71^. The unusual *S,S* – stereoconfiguration has been proposed to be responsible for the high stability of BMP in acidic organelles, because phospholipases preferentially hydrolyze phospholipids with *R-*configuration^70^. Based on the *R,S -* configuration of PG and LPG, head group acylation of these metabolites will initially result in products with *R,S -* configuration, which are subsequently converted into *S,S -* isomers. Consequently, all of the currently known PG/LPG transacylases likely produce *R,S -* precursors of *S,S -* BMP. The mechanism of stereoconversion is currently unclear. Our study suggests that BMP synthesis is not accompanied by a substantial exchange of glycerol moieties in the di(glycero)phosphate backbone. Thus, BMP synthesis does apparently not require the degradation and re-synthesis of the backbone of PG. We speculate that the enzymatic conversion of PG into BMP is accompanied by configurational changes leading to *S,S -* isomers, possibly via a cyclic intermediary product (**Figure 6B**). A similar mechanism was already proposed by Thornburg *et al.* in 1991^72^. Cyclic products as intermediates of enzymatic reactions are not unusual and have been detected in phospholipase C- and D-catalyzed reactions^73^. However, additional studies are required to elucidate in detail whether enzyme-catalyzed BMP synthesis is accompanied by stereoconformational changes or if *S,S -* BMP synthesis requires additional steps.

In summary, we demonstrate that BMP or precursors thereof are synthesized via functionally overlapping pathways in intra- and extracellular compartments. Our findings are highly relevant for the pathogenesis of lysosomal storage disorders^74^ as well as a range of other diseases such as viral infections^3^ and autoimmune disease^26^. Considering that BMP improves lysosomal function and autophagic flux^10,12,75,76^, the demonstrated efficient conversion of PG to BMP *in vivo* harbors a considerable therapeutic promise for the treatment of common neurodegenerative diseases, which are usually associated with lysosomal dysfunction^77^. We expect that further detailed investigation of the pathways reported herein will improve our understanding of ILV formation and endolysosomal cargo sorting in health and disease.

---

**Figure S1.**
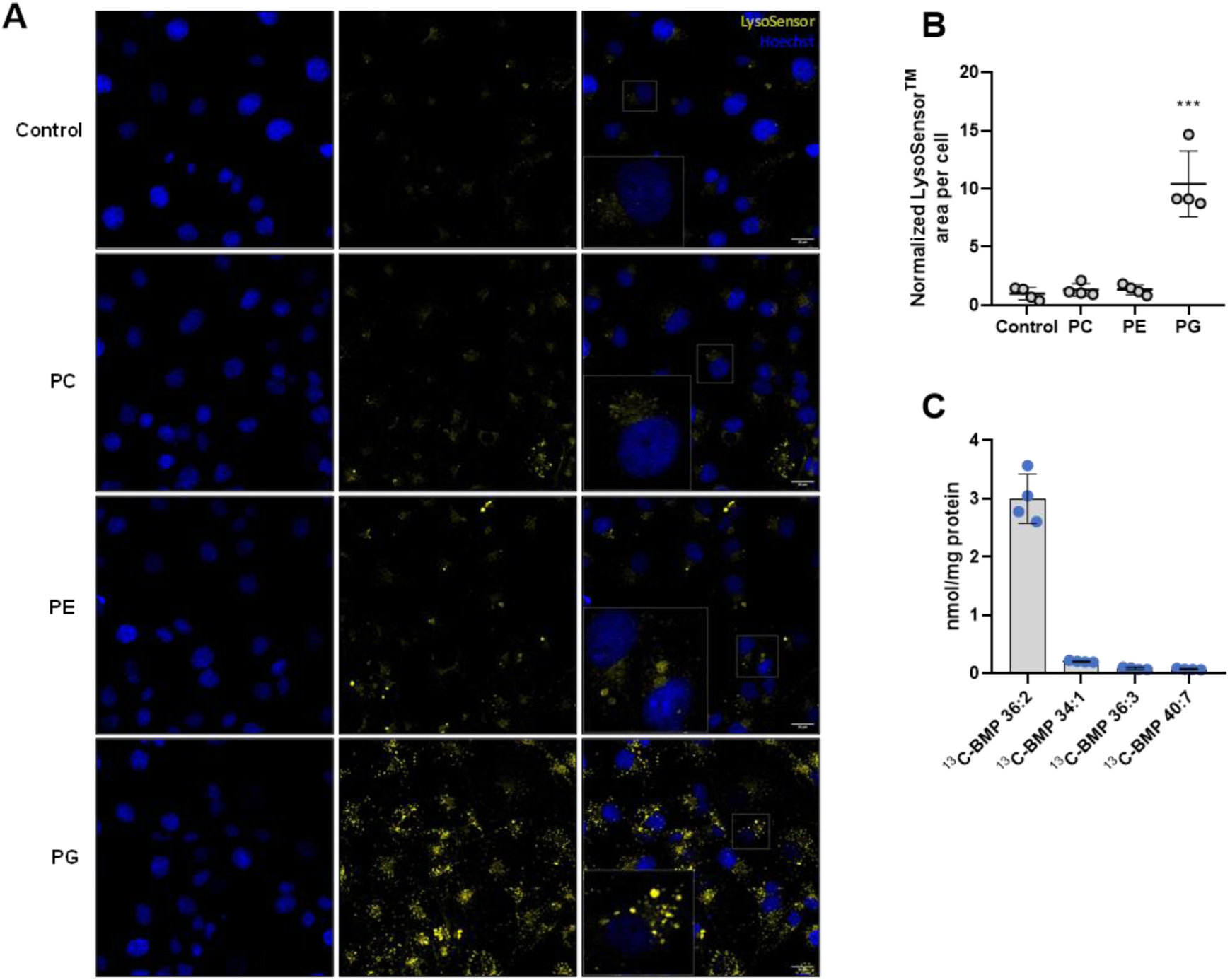
(**A**) Live cell imaging of COS-7 cells supplemented with indicated phospholipids (50 µM, all esterified with oleic acid). LE/Lysosomes were stained with LysoSensor^TM^ and nuclei were stained with Hoechst. (**B**) Quantification of LysoSensor^TM^ signal. The fluorescent signal of four randomly picked sections was determined in ImageJ and normalized to cell number (n=4). (**C**) ^13^C-labeled BMP subspecies formed during ^13^C-PG loading (n=4). Data are presented as mean ± SD. Statistically significant differences in (A) were determined by one-way ANOVA and corrected for multiple comparisons against control by Bonferroni post hoc test (levels of statistically significant differences are: ***p < 0.001).

**Figure S2.**
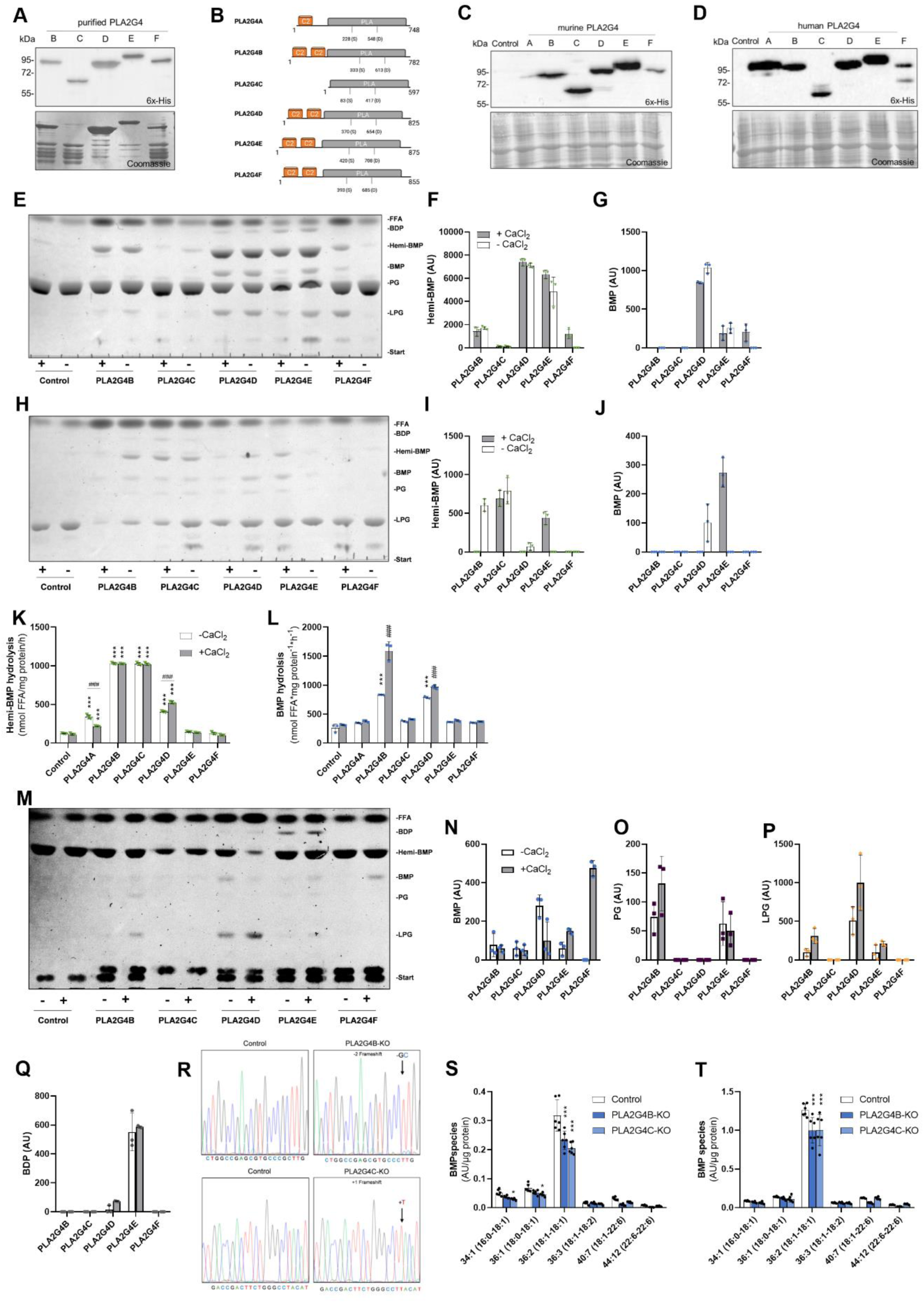
(**A**) Western blotting analysis of PLA2G4 enzymes enriched from lysates of Expi293F cells by affinity chromatography. Coomassie staining indicates that enzymes were partially purified. (**B**) Graphical illustrations of the domains and active sites of the murine PLA2G4 family. (**C, D**) Western blotting analysis of mouse and human PLA2G4A-F expression in Expi293F cells. The lower image shows the Coomassie-stained blot as loading control. (**E**) Representative TLC showing reaction products of semi-purified PLA2G4B-F incubated with di-oleoyl PG in the presence and absence of CaCl_2_, respectively. (**F, G**) Densitometric analysis of hemi-BMP and BMP formation shown in (E) (n=3). (**H**) Representative TLC showing reaction products of purified PLA2G4B-F incubated with 1-oleoyl LPG in the presence and absence of CaCl_2_. (**I, J**) Densitometric analysis of hemi-BMP and BMP formation shown in (H) (n=3). (**K, L**) Hydrolysis of tri-oleoyl hemi-BMP and di-oleoyl BMP without and with CaCl_2_ (n=3). Lysates of Expi293F cells expressing recombinant murine PLA2G4 enzymes were used as source of enzymatic activity. (**M**) Representative TLC showing the reaction products formed upon the hydrolysis of tri-oleoyl hemi-BMP using semi-purified PLA2G4 enzymes. (**N-Q**) Densitometric analysis of (M) (n=3). (**R**) Verification of the CRISPR-Cas9 mediated deletion of *PLA2G4B* and *PLA2G4C*. Chromatograms of control cells and KO cells are shown. Black arrow bars highlight the region where a frameshift was introduced. (**S, T**) BMP FA-composition of control, *PLA2G4B*-KO, and *PLA2G4C*-KO cells under basal conditions and after PG supplementation (n=6). The labeling on the x-axis indicates the FA composition of the respective BMP subspecies (number of carbon atoms: degree of saturation). Data are shown as mean ± SD. Statistically significant differences were determined by two-way ANOVA, followed by corrections for multiple comparisons (* = compared to the respective control, ^#^ = comparison between groups) using Bonferroni post hoc test (levels of statistically significant differences are: **,^##^*p* < 0.01, and ***,^###^*p* < 0.001).

**Figure S3.**
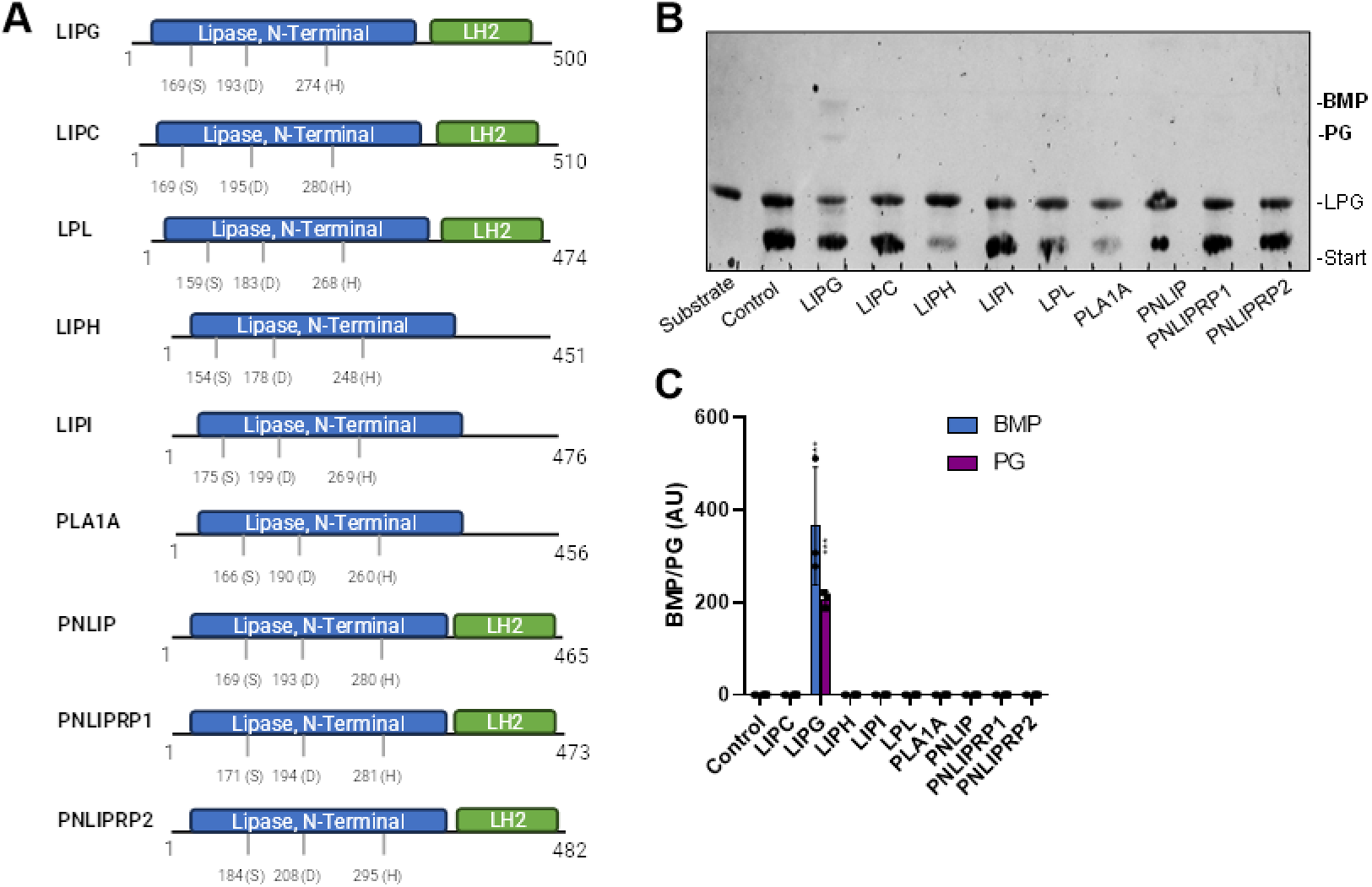
(**A**) Graphical illustrations of the domains and active sites of the murine PNLIP enzyme family. (**B**) Representative TLC showing LPG transacylase activity of PNLIP family members using 1-oleoyl LPG as substrate. (**C**) Densitometric analysis of BMP and PG formation shown in (B) (n=3). Data are presented as mean ± SD. Statistically significant differences were determined by two-way ANOVA, followed by corrections for multiple comparisons versus control using Bonferroni post hoc test (levels of statistically significant differences are: \*\*\**p* < 0.001).

**Figure S4.**
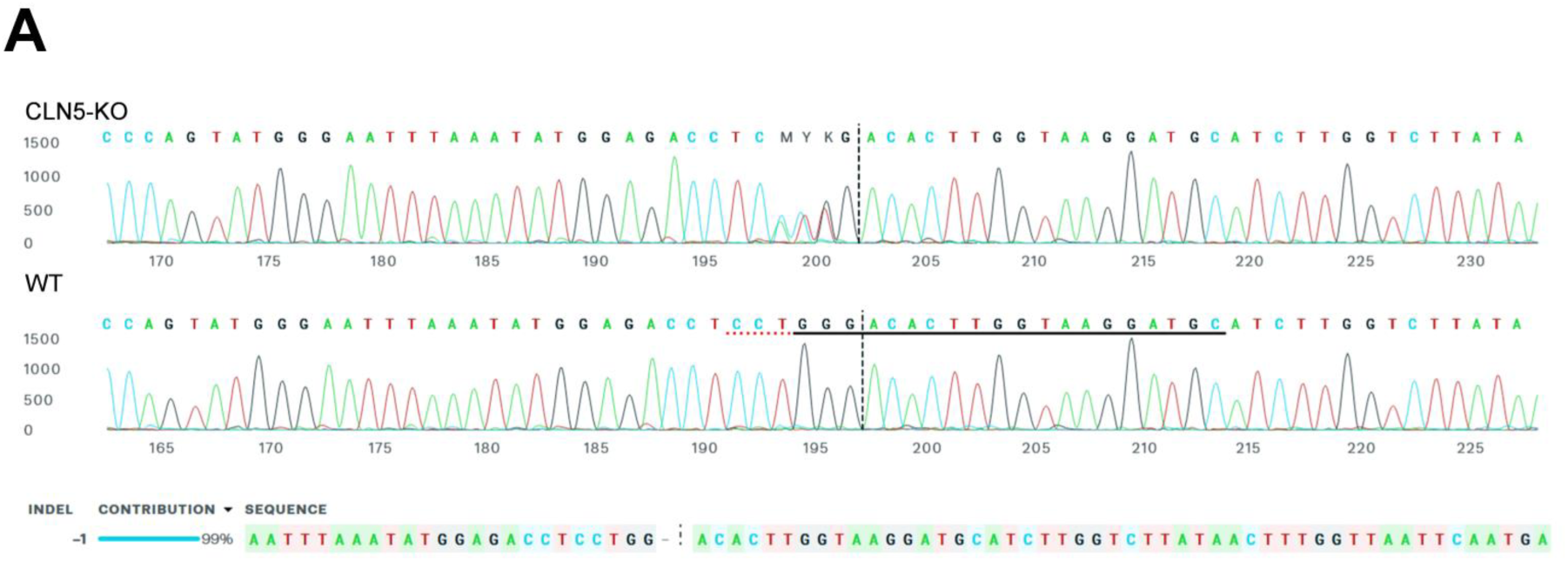
(A) Verification of CLN5 knockout in HEK293 cells by Sanger sequencing. The sequence was analyzed using the ICE CRISPR analysis tool by Synthego.

**Figure S5.**
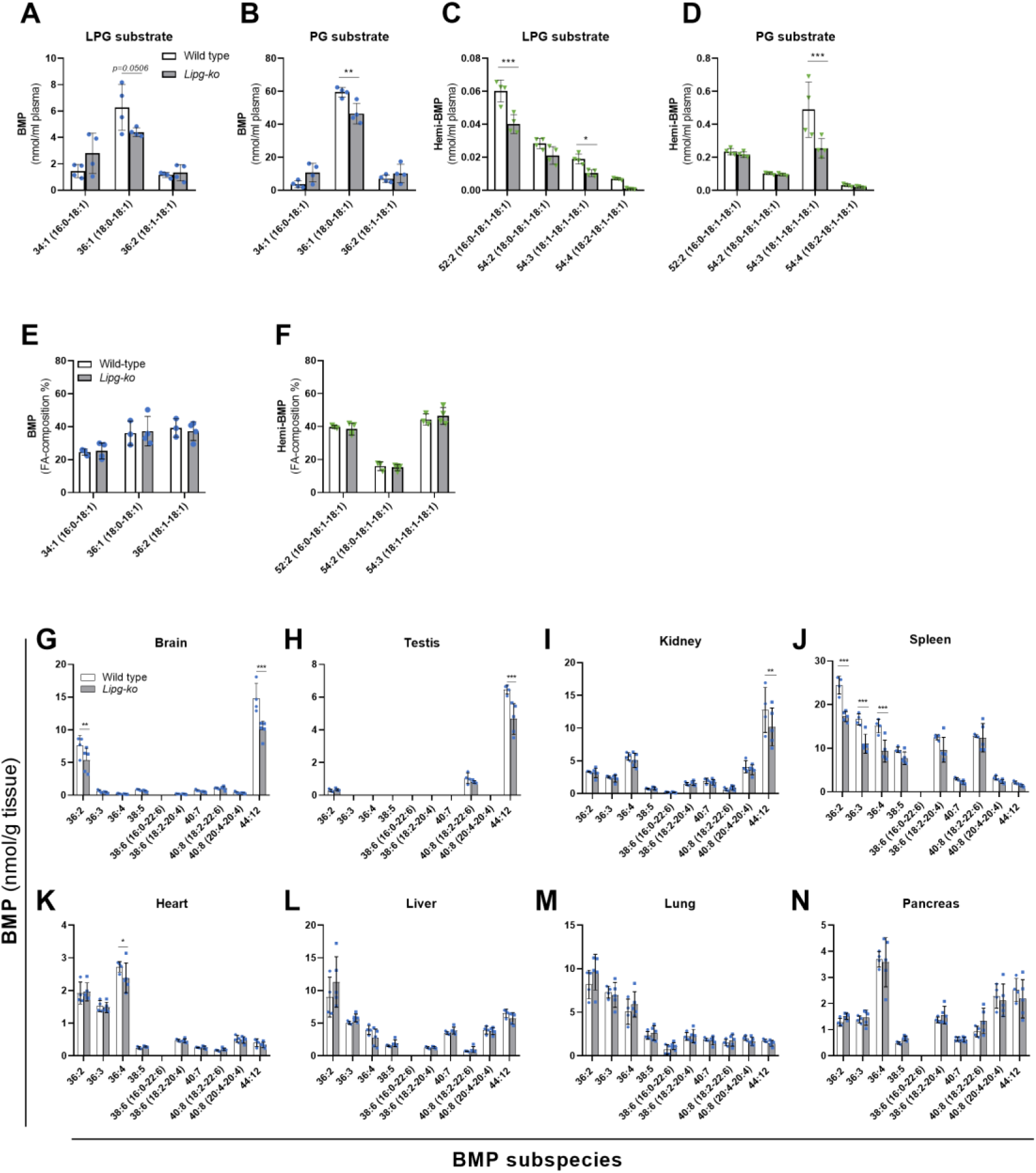
(**A, B**) BMP and (**C, D**) hemi-BMP subspecies formed in heparin-plasma of wild-type and *Lipg*-ko mice using oleoyl-LPG or di-oleoyl PG as substrate (n=4). (**E&F**) Relative composition of plasma BMP and hemi-BMP subspecies of wild-type and *Lipg*-ko mice (n=3 and 4, respectively). Blood samples were collected one hour after injection of di-oleoyl PG (5mg/mouse). (**G-N**) BMP composition of the indicated tissues (wild-type n=4, *Lipg*-ko n=5). The labeling on the x-axis indicates the FA composition of the respective BMP subspecies (number of carbon atoms: degree of saturation). Data are shown as mean ± SD. Statistically significant differences were evaluated by two-way ANOVA, followed by corrections for multiple comparisons using Bonferroni post hoc test (levels of statistically significant differences are: \**p* < 0.05, \*\**p* < 0.01, \*\*\**p* < 0.001).

## Data availability

Source data are provided with this paper as a Source Data file. All other data (i.e., validation of the screening library, uncropped TLCs, and Western blots) are deposited at Mendeley Data and openly accessible (DOI: 10.17632/hcs7crxk3z.1).

## Supplemental information

Supplemental information will we provided separately.

Supplemental Table S1: **List of qPCR primers, gRNAs, and sequencing primers**

Supplemental Table S2: **List of recombinant proteins and their respective vectors.**

## Acknowledgments

We want to thank Kathrin A. Zierler, Birgit Juritsch, and Sharon Fiorin for animal care and genotyping. We also thank R. Farese, T. Walther, and Shubham Singh (Sloan Kettering Institute, NY) for helpful discussions.

## Author contributions

Conceptualization, D.K., P.G., and R.Z.; Methodology, D.B., J.B., G.G., N.F., M.T., C.Z, A.P., G.B., H.W. and D.Ko.; Validation, L.H., A.L., and U.T., and N.C.; Formal Analysis; D.B., J.B., G.G., N.F., A.P., and D.Ko.; Investigation, D.B., J.B., G.G., N.F., M.T., C.Z, A.P., H.W. and D.Ko.; Writing – Original Draft, D.B., D.K., P.G., and R.Z.; Visualization, D.B., J.B., G.G., H.W. and D.Ko.; Supervision, D.K., A.L., and R.Z.

## REFERENCES

1. Huotari, J. & Helenius, A. Endosome maturation. EMBO J 30, 3481–3500 (2011).

2. Hullin-Matsuda, F., Taguchi, T., Greimel, P. & Kobayashi, T. Lipid compartmentalization in the endosome system. Semin. Cell Dev. Biol. 31, 48–56 (2014).

3. Gruenberg, J. Life in the lumen: The multivesicular endosome. Traffic 21, 76–93 (2020).

4. Hullin-Matsuda, F., Luquain-Costaz, C., Bouvier, J. & Delton-Vandenbroucke, I. Bis(monoacylglycero)phosphate, a peculiar phospholipid to control the fate of cholesterol: Implications in pathology. Prostaglandins Leukot Essent Fat. Acids 81, 313–324 (2009).

5. Gallala, H. D. & Sandhoff, K. Biological function of the cellular lipid BMP-BMP as a key activator for cholesterol sorting and membrane digestion. Neurochem Res 36, 1594– 1600 (2011).

6. Grumet, L. et al. Lysosomal Acid Lipase Hydrolyzes Retinyl Ester and Affects Retinoid Turnover. J. Biol. Chem. 291, 17977–17987 (2016).

7. Sandhoff, K. Metabolic and cellular bases of sphingolipidoses. Biochem. Soc. Trans. 41, 1562–1568 (2013).

8. Abdul-Hammed, M., al., et, Breiden, B., Schwarzmann, G. & Sandhoff, K. Lipids regulate the hydrolysis of membrane bound glucosylceramide by lysosomal β-glucocerebrosidase. J. Lipid Res. 58, 563–577 (2017).

9. Kobayashi, T. et al. Late endosomal membranes rich in lysobisphosphatidic acid regulate cholesterol transport. Nat. Cell Biol. 1, 113–8 (1999).

10. Chevallier, J. et al. Lysobisphosphatidic Acid Controls Endosomal Cholesterol Levels. J. Biol. Chem. 283, 27871–27880 (2008).

11. Xu, Z., Farver, W., Kodukula, S. & Storch, J. Regulation of sterol transport between membranes and NPC2. Biochemistry 47, 11134–11143 (2008).

12. McCauliff, L. A. et al. Intracellular cholesterol trafficking is dependent upon NPC2 interaction with lysobisphosphatidic acid. Elife 8, (2019).

13. Moreau, D. et al. Drug-induced increase in lysobisphosphatidic acid reduces the cholesterol overload in Niemann-Pick type C cells and mice. EMBO Rep. 20, (2019).

14. Meikle, P. J. et al. Effect of lysosomal storage on bis(monoacylglycero)phosphate. Biochem J 411, 71–78 (2008).

15. Reasor, M. J., Hastings, K. L. & Ulrich, R. G. Drug-induced phospholipidosis: issues and future directions. Expert Opin. Drug Saf. 5, 567–583 (2006).

16. Merchant, K. M. et al. LRRK2 and GBA1 variant carriers have higher urinary bis(monacylglycerol) phosphate concentrations in PPMI cohorts. NPJ Park. Dis. 9, (2023).

17. Chan, R. B. et al. Comparative lipidomic analysis of mouse and human brain with Alzheimer disease. J. Biol. Chem. 287, 2678–2688 (2012).

18. Miranda, A. M. et al. Effects of APOE4 allelic dosage on lipidomic signatures in the entorhinal cortex of aged mice. Transl. Psychiatry 12, (2022).

19. Akgoc, Z. et al. Bis(monoacylglycero)phosphate: a secondary storage lipid in the gangliosidoses. J. Lipid Res. 56, 1006–1013 (2015).

20. Boland, S. et al. Deficiency of the frontotemporal dementia gene GRN results in gangliosidosis. Nat. Commun. 13, (2022).

21. Logan, T. et al. Rescue of a lysosomal storage disorder caused by Grn loss of function with a brain penetrant progranulin biologic. Cell 184, 4651–4668.e25 (2021).

22. Medoh, U. N. et al. The Batten disease gene product CLN5 is the lysosomal bis(monoacylglycero)phosphate synthase. Science 381, 1182–1189 (2023).

23. Gardner, E. & Mole, S. E. The Genetic Basis of Phenotypic Heterogeneity in the Neuronal Ceroid Lipofuscinoses. Front. Neurol. 12, (2021).

24. Vreede, A. P., Bockenstedt, P. L. & Knight, J. S. Antiphospholipid syndrome. Curr. Opin. Rheumatol. 29, 458–466 (2017).

25. Vandevelde, A. & Devreese, K. M. J. Laboratory Diagnosis of Antiphospholipid Syndrome: Insights and Hindrances. J. Clin. Med. 11, 2164 (2022).

26. Müller-Calleja, N. et al. Lipid presentation by the protein C receptor links coagulation with autoimmunity. Science 371, (2021).

27. Le Blanc, I. et al. Endosome-to-cytosol transport of viral nucleocapsids. Nat Cell Biol 7, 653–664 (2005).

28. Mannsverk, S., Villamil Giraldo, A. M. & Kasson, P. M. Influenza Virus Membrane Fusion Is Promoted by the Endosome-Resident Phospholipid Bis(monoacylglycero)phosphate. J. Phys. Chem. B 126, (2022).

29. Luquain-Costaz, C., Rabia, M., Hullin-Matsuda, F. & Delton, I. Bis(monoacylglycero)phosphate, an important actor in the host endocytic machinery hijacked by SARS-CoV-2 and related viruses. Biochimie 179, 247–256 (2020).

30. van Blitterswijk, W. J. & Hilkmann, H. Rapid attenuation of receptor-induced diacylglycerol and phosphatidic acid by phospholipase D-mediated transphosphatidylation: formation of bisphosphatidic acid. EMBO J. 12, 2655–2662 (1993).

31. Hullin-Matsuda, F. et al. De novo biosynthesis of the late endosome lipid, bis(monoacylglycero)phosphate. J. Lipid Res. 48, 1997–2008 (2007).

32. Waite, M., King, L., Thornburg, T., Osthoff, G. & Thuren, T. Y. Metabolism of phosphatidylglycerol and bis(monoacylglycero)-phosphate in macrophage subcellular fractions. J Biol Chem 265, 21720–21726 (1990).

33. Labun, K. et al. CHOPCHOP v3: expanding the CRISPR web toolbox beyond genome editing. Nucleic Acids Res. 47, W171–W174 (2019).

34. du Sert, N. P., et al. Reporting animal research: Explanation and elaboration for the ARRIVE guidelines 2.0. PLOS Biol. 18, e3000411 (2020).

35. Juneja, L. R., Hibi, N., Inagaki, N., Yamane, T. & Shimizu, S. Comparative study on conversion of phosphatidylcholine to phosphatidylglycerol by cabbage phospholipase D in micelle and emulsion systems. Enzyme Microb. Technol. 9, 350–354 (1987).

36. Matyash, V., Liebisch, G., Kurzchalia, T. V., Shevchenko, A. & Schwudke, D. Lipid extraction by methyl-tert-butyl ether for high-throughput lipidomics. J. Lipid Res. 49, 1137–1146 (2008).

37. Folch, J., Lees, M. & Sloane Stanley, G. H. A simple method for the isolation and purification of total lipides from animal tissues. J. Biol. Chem. 226, 497–509 (1957).

38. Schindelin, J., et al. Fiji: an open-source platform for biological-image analysis. Nat. Methods 9, 676–682 (2012).

39. Bouvier, J. et al. Selective decrease of bis(monoacylglycero)phosphate content in macrophages by high supplementation with docosahexaenoic acid. J Lipid Res 50, 243–255 (2009).

40. Waite, M., Roddick, V., Thornburg, T., King, L. & Cochran, F. Conversion of phosphatidylglycerol to lyso(bis)phosphatidic acid by alveolar macrophages. FASEB J. 1, 318–325 (1987).

41. Ollis, D. L. & Carr, P. D. Hydrolase Fold: An Update. Protein Pept. Lett. 16, 1137– 1148 (2009).

42. Kienesberger, P. C., Oberer, M., Lass, A. & Zechner, R. Mammalian patatin domain containing proteins: A family with diverse lipolytic activities involved in multiple biological functions. Journal of Lipid Research vol. 50 at 10.1194/jlr.R800082-JLR200 (2009).

43. Mardian, E. B., Bradley, R. M. & Duncan, R. E. The HRASLS (PLA/AT) subfamily of enzymes. J. Biomed. Sci. 22, (2015).

44. Ghosh, M., Tucker, D. E., Burchett, S. A. & Leslie, C. C. Properties of the Group IV phospholipase A2 family. Prog. Lipid Res. 45, 487–510 (2006).

45. Hogan, S. et al. Studies on the antiobesity activity of tetrahydrolipstatin, a potent and selective inhibitor of pancreatic lipase. Int. J. Obes. 11 **Suppl 3**, 35–42 (1987).

46. Yasuda, T., Ishida, T. & Rader, D. J. Update on the role of endothelial lipase in high-density lipoprotein metabolism, reverse cholesterol transport, and atherosclerosis. Circ. J. 74, 2263–2270 (2010).

47. Lowe, M. E. The triglyceride lipases of the pancreas. J. Lipid Res. 43, 2007–16 (2002).

48. Broedl, U. C., Jin, W., Fuki, I. V., Glick, J. M. & Rader, D. J. Structural basis of endothelial lipase tropism for HDL. FASEB J. 18, 1891–1893 (2004).

49. Body, D. R. & Gray, G. M. The isolation and characterisation of phosphatidylglycerol and a structural isomer from pig lung. Chem. Phys. Lipids 1, 254–263 (1967).

50. Chen, J. et al. Lysosomal phospholipase A2 contributes to the biosynthesis of the atypical late endosome lipid bis(monoacylglycero)phosphate. *Commun*. Biol. 6, (2023).

51. Yamashita, A. et al. Subcellular localization and lysophospholipase/transacylation activities of human group IVC phospholipase A2 (cPLA2gamma). Biochim. Biophys. Acta 1791, 1011–1022 (2009).

52. Ogura, Y., Parsons, W. H., Kamat, S. S. & Cravatt, B. F. A calcium-dependent acyltransferase that produces N-acyl phosphatidylethanolamines. Nat. Chem. Biol. 12, 669–671 (2016).

53. Capestrano, M. et al. Cytosolic phospholipase A₂ε drives recycling through the clathrin-independent endocytic route. J. Cell Sci. 127, 977–993 (2014).

54. Yaginuma, S., Kawana, H. & Aoki, J. Current Knowledge on Mammalian Phospholipase A1, Brief History, Structures, Biochemical and Pathophysiological Roles. Molecules 27, (2022).

55. Young, S. G. & Zechner, R. Biochemistry and pathophysiology of intravascular and intracellular lipolysis. Genes and Development vol. 27 459–484 at 10.1101/gad.209296.112 (2013).

56. Record, M. et al. Bis (monoacylglycero) phosphate interfacial properties and lipolysis by pancreatic lipase-related protein 2, an enzyme present in THP-1 human monocytes. Biochim Biophys Acta 1811, 419–430 (2011).

57. Sindelar, P. & Valtersson, C. Hepatic lipase acylates dolichol in the presence of a plasma cofactor in vitro. Biochemistry 32, 9508–9512 (1993).

58. Borrelli, G. M. & Trono, D. Recombinant Lipases and Phospholipases and Their Use as Biocatalysts for Industrial Applications. Int. J. Mol. Sci. 16, 20774–20840 (2015).

59. Basak, I. et al. A lysosomal enigma CLN5 and its significance in understanding neuronal ceroid lipofuscinosis. Cell. Mol. Life Sci. 78, 4735–4763 (2021).

60. Furse, S. Is phosphatidylglycerol essential for terrestrial life? J. Chem. Biol. 10, (2016).

61. Grabner, G. F. et al. Metabolic disease and ABHD6 alter the circulating bis(monoacylglycerol)phosphate profile in mice and humans. J. Lipid Res. jlr.M093351 (2019) doi:10.1194/jlr.M093351.

62. Meikle, P., Duplock, S. & Blacklock, D. Effect of lysosomal storage on bis (monoacylglycero) phosphate. Biochem. J 411, 71–78 (2008).

63. Rabia, M. et al. Bis(monoacylglycero)phosphate, a new lipid signature of endosome-derived extracellular vesicles. Biochimie 178, 26–38 (2020).

64. Matsuzawa, Y. & Hostetler, K. Y. Degradation of bis(monoacylglycero)phosphate by an acid phosphodiesterase in rat liver lysosomes. J Biol Chem 254, 5997–6001 (1979).

65. Pribasnig, M. A. et al. α/β Hydrolase Domain-containing 6 (ABHD6) Degrades the Late Endosomal/Lysosomal Lipid Bis(monoacylglycero)phosphate. J. Biol. Chem. 290, 29869–81 (2015).

66. Aoki, J., Inoue, A., Makide, K., Saiki, N. & Arai, H. Structure and function of extracellular phospholipase A1 belonging to the pancreatic lipase gene family. Biochimie 89, 197–204 (2007).

67. Grabner, G. F. et al. Metabolic regulation of the lysosomal cofactor bis(monoacylglycero)phosphate in mice. J. Lipid Res. 61, (2020).

68. Silva, B. De, Adams, J. & Lee, S. Y. Proteolytic processing of the neuronal ceroid lipofuscinosis related lysosomal protein CLN5. Exp. Cell Res. 338, 45–53 (2015).

69. Holmberg, V. et al. The mouse ortholog of the neuronal ceroid lipofuscinosis CLN5 gene encodes a soluble lysosomal glycoprotein expressed in the developing brain. Neurobiol. Dis. 16, 29–40 (2004).

70. Tan, H. H., Makino, A., Sudesh, K., Greimel, P. & Kobayashi, T. Spectroscopic evidence for the unusual stereochemical configuration of an endosome-specific lipid. Angew Chem Int Ed Engl 51, 533–535 (2012).

71. Kobayashi, T. et al. Separation and Characterization of Late Endosomal Membrane Domains. J. Biol. Chem. 277, 32157–32164 (2002).

72. Thornburg, T., Miller, C., Thuren, T., King, L. & Waite, M. Glycerol reorientation during the conversion of phosphatidylglycerol to bis(monoacylglycerol)phosphate in macrophage-like RAW 264.7 cells. J. Biol. Chem. 266, 6834–6840 (1991).

73. Friedman, P., Haimovitz, R., Markman, O., Roberts, M. F. & Shinitzky, M. Conversion of lysophospholipids to cyclic lysophosphatidic acid by phospholipase D. J. Biol. Chem. 271, 953–957 (1996).

74. Showalter, M. R. et al. The Emerging and Diverse Roles of Bis(monoacylglycero) Phosphate Lipids in Cellular Physiology and Disease. Int. J. Mol. Sci. 21, 1–19 (2020).

75. Ilnytska, O. et al. Enrichment of NPC1-deficient cells with the lipid LBPA stimulates autophagy, improves lysosomal function, and reduces cholesterol storage. J. Biol. Chem. 297, (2021).

76. Ilnytska, O. et al. Lysobisphosphatidic acid (LBPA) enrichment promotes cholesterol egress via exosomes in Niemann Pick type C1 deficient cells. Biochim. Biophys. acta. Mol. cell Biol. lipids 1866, 158916 (2021).

77. Udayar, V., Chen, Y., Sidransky, E. & Jagasia, R. Lysosomal dysfunction in neurodegeneration: emerging concepts and methods. Trends Neurosci. 45, 184–199 (2022).

